# Systems biology informed deep learning for inferring parameters and hidden dynamics

**DOI:** 10.1101/865063

**Authors:** Alireza Yazdani, Lu Lu, Maziar Raissi, George Em Karniadakis

**Affiliations:** Division of Applied Mathematics, Brown University, Providence, RI 02912, USA; Department of Applied Mathematics, University of Colorado, Boulder, CO 80309, USA

## Abstract

Mathematical models of biological reactions at the system-level lead to a set of ordinary differential equations with many unknown parameters that need to be inferred using relatively few experimental measurements. Having a reliable and robust algorithm for parameter inference and prediction of the hidden dynamics has been one of the core subjects in systems biology, and is the focus of this study. We have developed a new systems-biology-informed deep learning algorithm that incorporates the system of ordinary differential equations into the neural networks. Enforcing these equations effectively adds constraints to the optimization procedure that manifests itself as an imposed structure on the observational data. Using few scattered and noisy measurements, we are able to infer the dynamics of unobserved species, external forcing, and the unknown model parameters. We have successfully tested the algorithm for three different benchmark problems.

**Author summary:** The dynamics of systems biological processes are usually modeled using ordinary differential equations (ODEs), which introduce various unknown parameters that need to be estimated efficiently from noisy measurements of concentration for a few species only. In this work, we present a new “systems-informed neural network” to infer the dynamics of experimentally unobserved species as well as the unknown parameters in the system of equations. By incorporating the system of ODEs into the neural networks, we effectively add constraints to the optimization algorithm, which makes the method robust to noisy and sparse measurements.

## Introduction

Systems biology aims at a system-level understanding of biological systems, which is a holistic approach to deciphering the complexity of biological systems [1]. To understand the biological systems, we must understand the structures of the systems (both their components and structural relationships), and their dynamics [2]. The dynamics of systems biological processes are usually modeled using ordinary differential equations (ODEs) that describe the time evolution of chemical and molecular species concentrations. Once the pathway structure of chemical reactions is known, the corresponding equations can be derived using widely accepted kinetic laws, such as the law of mass action or the Michaelis-Menten kinetics [3].

Most system-level biological models introduce various unknown parameters, which need to be estimated efficiently. Thus, the central challenge in computational modeling of these systems could be the prediction of model parameters such as rate constants or initial concentrations, and model trajectories such as time evolution of experimentally unobserved concentrations. Due to the importance of parameter estimation, a lot of attention has been given to this problem in the systems biology community. A lot of research has been conducted on the applications of several optimization techniques, such as linear and nonlinear least-squares fitting [4], genetic algorithms [5], evolutionary computation [6], and more [7]. Considerable interest has also been raised by Bayesian methods [8, 9], which can extract information from noisy or uncertain data. The main advantage of these methods is their ability to infer the whole probability distributions of the parameters, rather than just a point estimate. More recently, parameter estimation for computational biology models has been tackled in the framework of control theory by using state observers. These algorithms were originally developed for the problem of state estimation in which one seeks to estimate the time evolution of the unobserved components of the state of a dynamical system. In this context, extended Kalman filtering [10], unscented Kalman filtering [11], and ensemble Kalman methods [12] have been applied as well. In addition, different methods have also been developed to address the issue of hidden variables and dynamics [13, 14].

Due to technical limitations, however, biological reaction networks are often only partially observable. Usually, experimental data are insufficient considering the size of the model, which results in parameters that are non-identifiable [15] or only identifiable within confidence intervals (see more details in [16]). Furthermore, it is known that a large class of systems biology models display sensitivities to the parameter values that are roughly evenly distributed over many orders of magnitude. Such *sloppiness* has been suggested as a factor that makes parameter estimation difficult [17]. In the process of parameter inference, two issues accounting for system’s (non-)identifiability have to be considered: *structural* identifiability that is related to the model structure independent of the experimental data [18, 19]; and *practical* identifiability that takes into account the amount and quality of measured data. The *a priori* structural identifiability addresses the question of unique estimation of the unknown parameters based on the postulated model. However, a parameter that is structurally identifiable may still be practically non-identifiable assuming that the model is exact, but the measurements are noisy or sparse [20].

In this work, we introduce a new deep learning [21] method — systems-informed neural networks, based on the method of physics-informed neural networks [22, 23], to infer the *hidden* dynamics of experimentally unobserved species as well as the unknown parameters in the system of equations. By incorporating the system of ODEs into the neural networks (through adding the residuals of the equations to the loss function), we effectively add constraints to the optimization algorithm, which makes the method robust to measurement noise and few scattered observations. In addition, since large system-level biological models are typically encountered, our algorithm is computationally scalable and feasible, and its output is interpretable even though it depends on a high-dimensional parameter space.

## Materials and methods

Throughout this paper, we assume that the systems biological process can be modeled by a system of ordinary differential equations (ODEs) of the following form

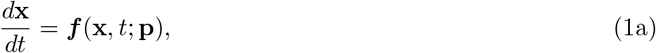

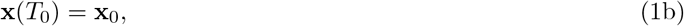

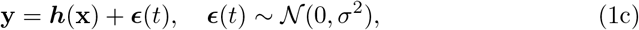

where the state vector **x** = (*x*_1_, *x*_2_, …, *x_S_*) represents the concentration of *S* species, and **p** = (*p*_1_, *p*_2_, …. *p_K_*) are *K* parameters of the model, which remain to be determined. Hence, the system of ODEs will be identified once **p** is known. **y** = (*y*_1_, *y*_2_, …, *y_M_*) are the *M* measurable signals (consistent with the ODE system), which we can measure experimentally and could possibly be contaminated with a noise ***ϵ*** of Gaussian type with zero mean and standard deviation *σ*. The output function ***h*** is determined by the design of the experiments that are used for parameter inference. While ***h*** could, in general, be any function, it is assumed to be a linear function with *M* ≤ *S* in most models as follows:

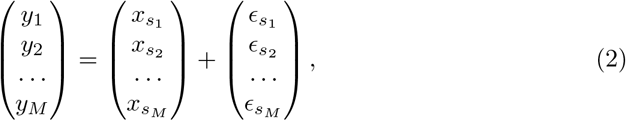

i.e., *y*_1_, *y*_2_, …, *y_M_* are the noisy measurements of the species *x*_*s*_1__, *x*_*s*_2__, …, *x_s_M__* among all *S* species (1 ≤ *s*_1_ < *s*_2_ < ⋯ < *s_M_* ≤ *S*).

### Systems-informed neural networks and parameter inference

Based on the method of physics-informed neural networks proposed in [22], we introduce a deep learning framework that is informed by the systems biology equations that describe the kinetic pathways (Eq. (1)a). A neural network with parameters ***θ*** takes time *t* as the input and outputs a vector of the state variables 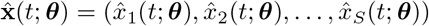 as a surrogate of the ODE solution **x**(*t*) (Fig. 1). In our network, in addition to the standard layers, e.g., the fully-connected layer, we add three extra layers to make the network training easier, described as follows.

- Input-scaling layer. Because the ODE system could have a large time domain, i.e., the input time t could vary by orders of magnitude, we first apply a linear scaling function to t, i.e., 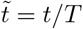, such that 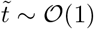. We can simply choose *T* as the maximum value of the time domain.
- Feature layer. In many models, the ODE solution may have a certain pattern, e.g., periodicity in the yeast glycolysis model and fast decay in the cell apoptosis model, and thus it is beneficial to the network training by constructing a feature layer according to these patterns. Specifically, we employ the *L* function *e*_1_(·), *e*_2_(·), …, *e_L_*(·) to construct the *L* features 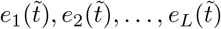.
- Output-scaling layer. Because the outputs 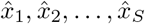 may have different magnitudes, similar to the input-scaling layer, we add another scaling layer to transform the output of the last fully-connected layer 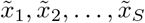 (of order one) to 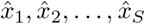, i.e., 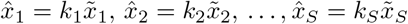. Here, *k*_1_, *k*_2_, …, *k_S_* are chosen as the magnitudes of the mean values of the ODE solution *x*_1_, *x*_2_, …, *x_S_*, respectively.

**Fig. 1.**
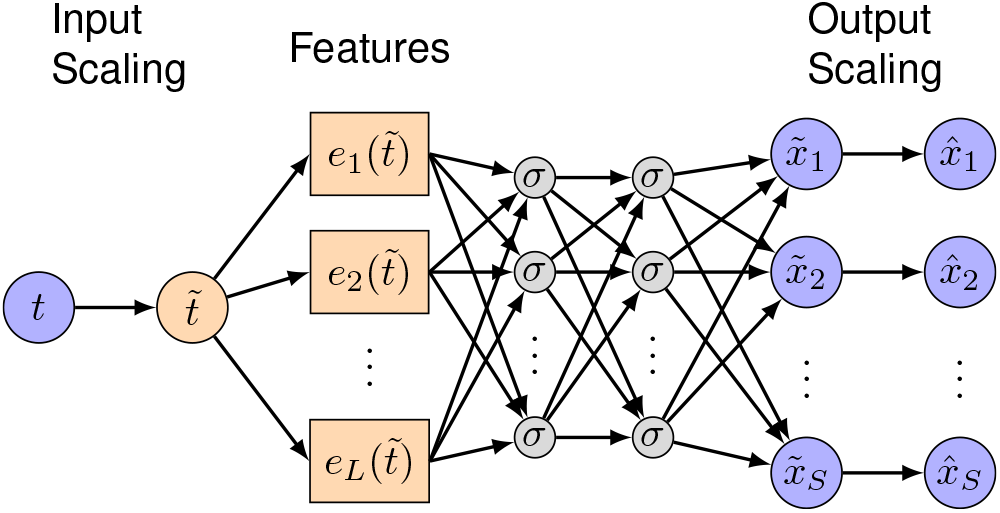
Neural network architecture. The network consists of an input-scaling layer, a feature layer, several fully-connected layers, and an output-scaling layer. The input-scaling layer and output-scaling layer are used to linearly scale the network input and outputs such that they are of order one. The feature layer is used to construct features explicitly as the input to the first fully-connected layer.

The next key step is to constrain the neural network to satisfy the scattered observations of **y** as well as the ODE system (Eq. (1)a). This is realized by constructing the loss function by considering terms corresponding to the observations and the ODE system. Specifically, let us assume that we have the measurements of *y*_1_, *y*_2_, …, *y_M_* at the time *t*_1_, *t*_2_, …, *t_N^data^_*, and we enforce the network to satisfy the ODE system at the time point *τ*_1_, *τ*_2_, …, *τ_N^ode^_*. We note that the times *t*_1_, *t*_2_, …, *t_N^data^_* and *τ*_1_, *τ*_2_, …, *τ_N^ode^_* are not necessarily on a uniform grid, and they could be chosen at random. Then, the total loss is defined as a function of both ***θ*** and **p**:

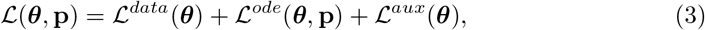

where

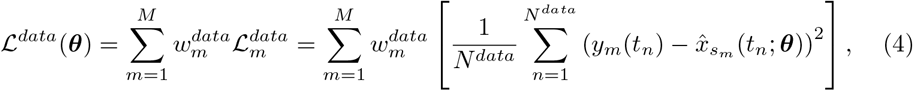

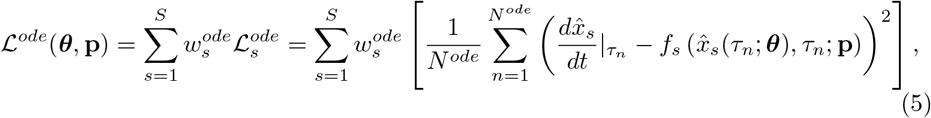

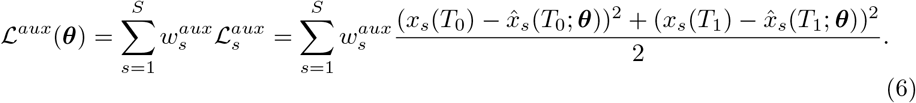

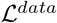 is associated with the *M* sets of observations **y** given by Eq. (1)c, while 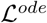 enforces the structure imposed by the system of ODEs given in Eq. (1)a. We employ automatic differentiation (AD) to analytically compute the derivative 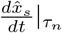 in 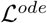 (see more details of AD in [23]). The third auxiliary loss term 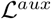 is introduced as an additional source of information for the system identification, and involves two time instants *T*_0_ and *T*_1_. It is essentially a component of the data loss; however, we prefer to separate this loss from the data loss, as in the auxiliary loss data are given for all state variables at these two time instants. We note that 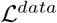 and 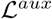 are the discrepancy between the network and measurements, and thus they are *supervised* losses, while 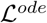 is based on the ODE system, and thus is *unsupervised*. In the last step, we infer the neural network parameters ***θ*** as well as the unknown parameters of the ODEs **p** simultaneously by minimizing the loss function via gradient-based optimizers, such as the Adam optimizer [24]:

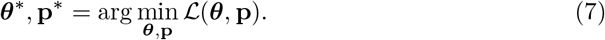

We note that our proposed method is different from meta-modeling [25, 26], as we optimize ***θ*** and **p** simultaneously.

The *M* + 2*S* coefficients 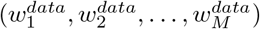 in Eq. (4), 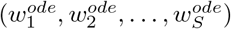 in Eq. (5), and 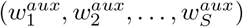 in Eq. (6) are used to balance the *M* + 2*S* loss terms. In this study, we manually select these weight coefficients such that the weighted losses are of the same order of magnitude during the network training. We note that this guideline makes the weight selection much easier, although there are many weights to be determined. These weights may also be automatically chosen, e.g., by the method proposed in [27]. In this study, the time instants *t*_1_, *t*_2_, …, *t_N^data^_* for the observations are chosen randomly in the time domain, while the time instants *τ*_1_, *τ*_2_, …, *τ_N^ode^_* used to enforce the ODEs are chosen in an equispaced grid. Additionally, in the auxiliary loss function, the first set of data is the initial conditions at time *T*_0_ for the state variables. The second set includes the values of the state variables at any arbitrary time instant *T*_1_ within the training time window (not too close to *T*_0_); in this work, we consider the midpoint time for the cell apoptosis model, and the final time instant for the yeast glycolysis model and ultradian endocrine model. If data is available at another time point, alternatively this point can be considered.

### Analysis of system’s identifiability

A parameter *p_i_* is identifiable if the confidence interval of its estimate 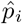 is finite. In systems identification problems, two different forms of identifiability namely, *structural* and *practical* are typically encountered. Structural non-identifiability arises from a redundant parameterization in the formal solution of **y** due to insufficient mapping **h** of internal states **x** to observables **y** in Eq. (1) [15]. *A priori* structural identifiability has been studied extensively, e.g., using power series expansion [28] and differential algebraic methods [29], yet mostly limited to linear models as the problem is particularly difficult for nonlinear dynamical systems. Furthermore, practical non-identifiability cannot be detected with these methods, since experimental data are disregarded.

A parameter that is structurally identifiable may still be practically non-identifiable. Practical non-identifiability is intimately related to the amount and quality of measured data and manifests in a confidence interval that is infinite. Different methods have been proposed to estimate the confidence intervals of the parameters such as local approximation of the Fisher-Information-Matrix (FIM) [20] and bootstrapping approach [30]. Another approach is to quantify the sensitivity of the systems dynamics to variations in its parameters using a probabilistic framework [31]. For identifiability analysis, we primarily use the FIM method, which is detailed in the Supporting Information.

### Implementation

The algorithm is implemented in Python using the open-source library DeepXDE [23]. The width and depth of the neural networks (listed in Table 1) depend on the size of the system of equations and the complexity of the dynamics. We use the 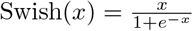 function [32] as the activation function *σ* shown in Fig. 1, and the feature layer is listed in Table **2**. The source codes for these three problems are available to download at https://github.com/alirezayazdani1/SBINNs.

**Table 1.** Hyperparameters for the problems in this study. The fist and second number in the number of iterations correspond to the first and second training stage.

| Model |  | NN depth | NN width | #Iterations |
| --- | --- | --- | --- | --- |
| Yeast glycolysis | Noiseless | 4 | 128 | 1000, $9 \times 10^4$ |
| | Noisy | 4 | 128 | 1000, $2 \times 10^6$ |
| Cell apoptosis | Survival | 5 | 256 | 0, $1.5 \times 10^6$ |
| | Death | 5 | 256 | 0, $1.5 \times 10^6$ |
| Ultradian endocrine | Parameters only | 4 | 128 | 2000, $6 \times 10^5$ |
| | Hidden nutrition | 4 | 128 | 2000, $1.5 \times 10^6$ |

**Table 2.** The feature layer used in the network for each problem.

| Model | Features |
| --- | --- |
| Yeast glycolysis | $\tilde{t}, \sin(\tilde{t}), \sin(2\tilde{t}), \sin(3\tilde{t}), \sin(4\tilde{t}), \sin(5\tilde{t}), \sin(6\tilde{t})$ |
| Cell apoptosis | $\tilde{t}, e^{-\tilde{t}}$ |
| Ultradian endocrine | $\tilde{t}, \sin(\tilde{t}), \sin(2\tilde{t}), \sin(3\tilde{t}), \sin(4\tilde{t}), \sin(5\tilde{t})$ |

For the training, we use an Adam optimizer [24] with default hyperparameters and a learning rate of 10^−3^, where the training is performed using the full batch of data. As the total loss consists of two supervised losses and one unsupervised loss, we perform the training using the following two-stage strategy:

Step 1 Considering that supervised training is usually easier than unsupervised training, we first train the network using the two supervised losses 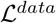 and 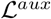 for some iterations, such that the network can quickly match the observed data points.
Step 2 We further train the network using all the three losses.

We found empirically that this two-stage training strategy speeds up the network convergence. The number of iterations for each stage is listed in Table **1**.

## Results

### Yeast glycolysis model

The model of oscillations in yeast glycolysis [33] has become a standard benchmark problem for systems biology inference [34, 35] as it represents complex nonlinear dynamics typical of biological systems. We use it here to study the performance of our deep learning algorithm used for parsimonious parameter inference with only two observables. The system of ODEs for this model as well as the target parameter values and the initial conditions are included in the Supporting Information. To represent experimental noise, we corrupt the observation data by a Gaussian noise with zero mean and the standard deviation of *σ_ϵ_* = *cμ*, where *μ* is the standard deviation of each observable over the observation time window and *c* = 0 – 0.1.

We start by inferring the dynamics using noiseless observations on two species *S*_5_ (the concentration of NADH) and *S*_6_ (the concentration of ATP) only. These two species are the minimum number of observables we can use to effectively infer all the parameters in the model. Fig. **S1** shows the noiseless synthetically generated data by solving the system of ODEs in Eq. (S4). We sample data points within the time frame of 0 – 10 minutes at random and use them for training of the neural networks, where the neural network is informed by the governing ODEs of the yeast model as explained above. Fig. **S2** shows the inferred dynamics for all the species predicted by the systems-informed neural networks, and plotted against the exact dynamics that are generated by solving the system of ODEs. We observe excellent agreement between the inferred and exact dynamics within the training time window. The neural networks learn the input data given by scattered observations (shown by symbols in Fig. S2) and is able to infer the dynamics of other species due to the constraints imposed by the system of ODEs.

Next, we verify the robustness of the algorithm to noise. For that purpose, we introduce Gaussian additive noise with the noise level *c* = 10% to the observational data. The input training data are shown in Fig. **2** for the same species (*S_5_* and *S_6_*) as the observables, where similar to the previous test, we sample random scattered data points in time. Results for the inferred dynamics are shown in Fig. **3**. The agreement between the inferred and exact dynamics is excellent considering the relatively high level of noise in the training data. Interestingly, our results show that the enforced equations in the loss function 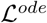 act as a constraint of the neural networks that can effectively prevent the overfitting of the network to the noisy data. One advantage of encoding the equations is their regularization effect without using any additional *L*_1_ or *L*_2_ regularization.

**Fig. 2.**
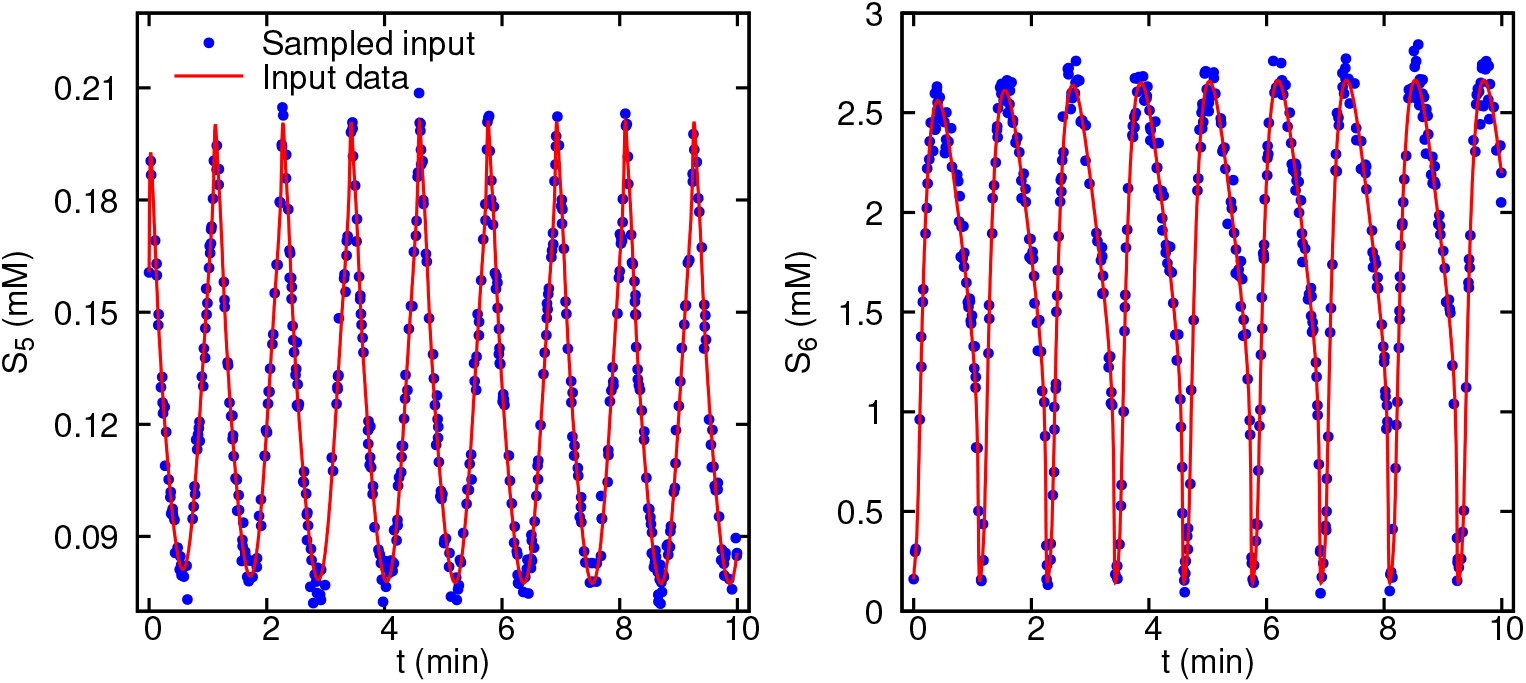
Glycolysis oscillator noisy observation data given to the algorithm for parameter inference. 500 measurements are corrupted by a zero-mean Gaussian noise and standard deviation of *σ* = 0.1*μ*. Only two observables *S*_5_ and *S*_6_ are considered and the data are randomly sampled in the time window of 0 – 10 minutes.

**Fig. 3.**
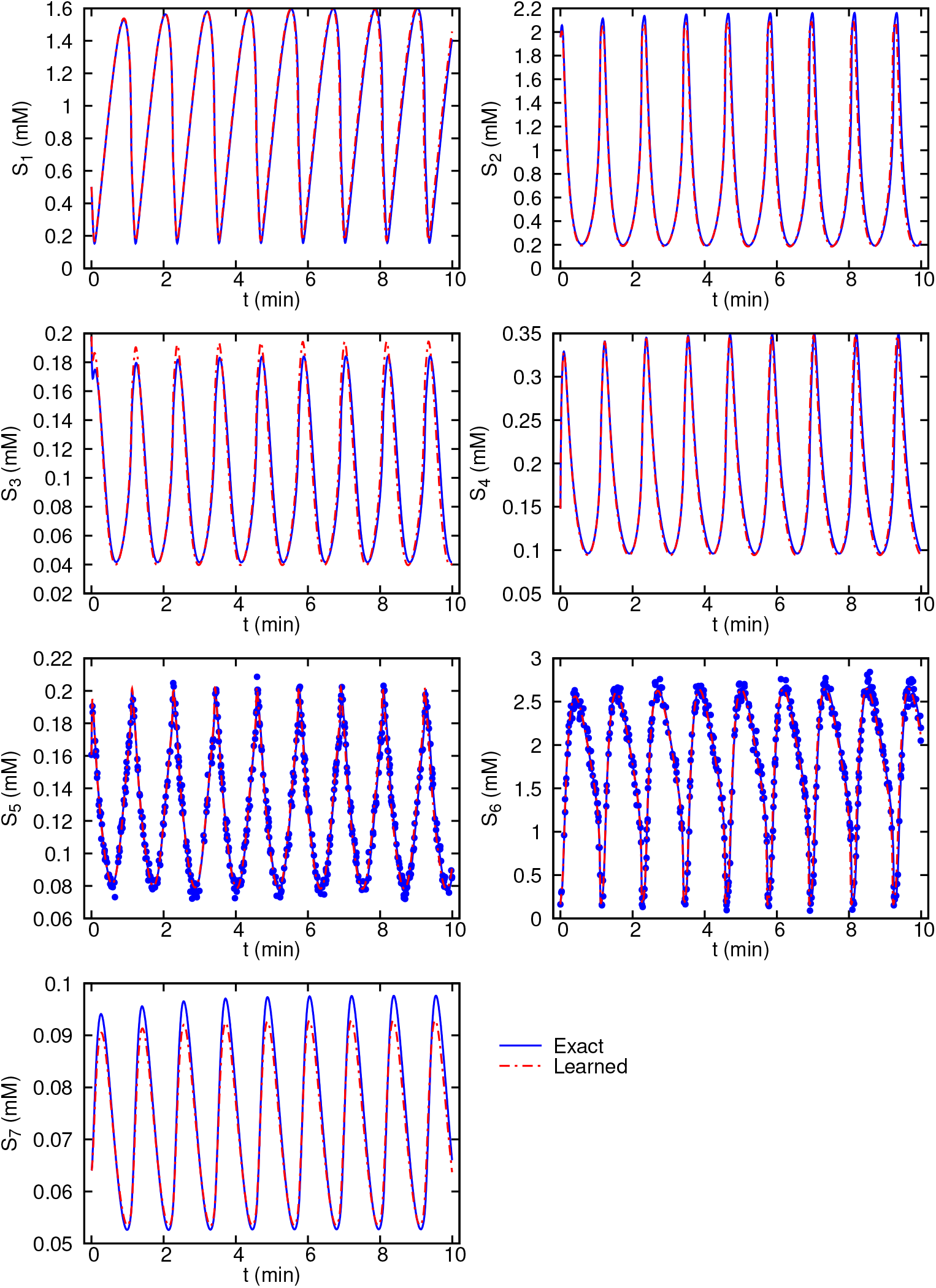
Glycolysis oscillator inferred dynamics from noisy measurements compared with the exact solution. 500 scattered observations are plotted using symbols for the two observables *S*_5_ and *S*_6_.

Our main objective in this work, however, is to infer the unknown model parameters **p**. This can be achieved simply by training the neural networks for its parameters ***θ*** as well as the model parameters using backpropagation. The results for the inferred model parameters along with their target values are given in Table **3** for both test cases (i.e., with and without noise in the observations). First thing to note is that the parameters can be identified within a confidence interval. Estimation of the confidence intervals *a priori* is the subject of structural identifiability analysis, which is not in the scope of this work. Second, practical identifiability analysis can be performed to identify the practically non-identifiable parameters based on the quality of the measurements and the level of the noise. We have performed local sensitivity analysis by constructing the Fisher Information Matrix (FIM) using Eq. (S1) and the correlation matrix **R** derived from the FIM.

**Table 3.** Parameter values for yeast glycolysis model and each corresponding inferred values. The standard deviations are estimated using Eq. (S3) as practical non-identifiability analysis based on the FIM.

| Parameter | Target value | Inferred value<br>(Noiseless observations) | Inferred value<br>(Noisy observations) | Standard<br>deviation |
| --- | --- | --- | --- | --- |
| $J_0$ | 2.5 | 2.50 | 2.49 | 0.18 |
| $k_1$ | 100 | 99.9 | 86.1 | 62.0 |
| $k_2$ | 6 | 6.01 | 4.55 | 21.3 |
| $k_3$ | 16 | 15.9 | 14.0 | 21.9 |
| $k_4$ | 100 | 100.1 | 97.1 | 103.6 |
| $k_5$ | 1.28 | 1.28 | 1.24 | 0.25 |
| $k_6$ | 12 | 12.0 | 12.7 | 5.1 |
| $k$ | 1.8 | 1.79 | 1.55 | 4.34 |
| $\kappa$ | 13 | 13.0 | 13.4 | 25.9 |
| $q$ | 4 | 4.00 | 4.07 | 0.27 |
| $K_1$ | 0.52 | 0.520 | 0.550 | 0.091 |
| $\psi$ | 0.1 | 0.0994 | 0.0823 | 0.317 |
| $N$ | 1 | 0.999 | 1.29 | 2.94 |
| $A$ | 4 | 4.01 | 4.25 | 2.28 |

The inferred parameters from both noiseless and noisy observations are in good agreement with their target values. The most significant difference can be seen for the parameter *N* (close to 30% difference). However, given that the glycolysis system of Eq. (S4) is identifiable (c.f. [33, 35] and Fig. S3), and the inferred dynamics shown in Figs. **S2** and **3** show that the learned dynamics match very well with the exact dynamics, the inferred parameters are valid. We used Eq. (S3) to estimate the standard deviations of the model parameters. The *σ_i_* estimates for the parameters are the lower bounds, and thus, may not be informative here. Further, these estimates are derived based on a local sensitivity analysis. A structural/practical identifiability analysis [15] or a bootstrapping approach to obtain the parameter confidence intervals is probably more relevant here. Using the FIM, we are able to construct the correlation matrix **R** for the parameters. Nearly perfect correlations (|*R_ij_*| ≈ 1) suggest that the FIM is singular and the correlated parameters may not be practically identifiable. For the glycolysis model, as shown in Fig. S3, no perfect correlations can be found in **R** (except for the anti-diagonal elements), which suggests that the model described by Eq. (S4) is practically identifiable. In the example above, we considered 500 data measurements, but in systems biology we often lack the ability to observe system dynamics at a fine-time scale. To investigate the performance of our method to a set of sparse data points, we used only 200 data points, and still have a good inferred dynamics of the species *S*_1_, *S*_2_, *S*_5_ and *S*_6_ (Fig. S4).

### Cell apoptosis model

Although the glycolysis model is highly nonlinear and difficult to learn, we have shown that its parameters can be identified. To investigate the performance of our algorithm for non-identifiable systems, we study a cell apoptosis model, which is a core sub-network of the signal transduction cascade regulating the programmed cell death-against-survival phenotypic decision [36]. The equations defining the cell apoptosis model and the values of the rate constants for the model are taken from [36] and listed in Table **S2** in the Supporting Information.

Although the model is derived using simple mass-action kinetics and its dynamics is easy to learn with our algorithm, most of the parameters are not identifiable due to both structural and practical non-identifiability. To infer the dynamics of this model, we only use 120 random samples of measurements collected for one observable (*x*_4_), where we assume that the measurements are corrupted by a zero-mean Gaussian noise and 5% standard deviation as shown in Fig. **4**. Furthermore, it is possible to use different initial conditions in order to produce different cell survival outcomes. The initial conditions for all the species are given in the Supporting Information, while we use *x*_7_(0) = 2.9 × 10^4^ (molecules/cell) to model cell survival (Fig. **4**(top)) and *x*_7_(0) = 2.9 × 10^3^ (molecules/cell) to model cell death (Fig. **4**(bottom)).

**Fig. 4.**
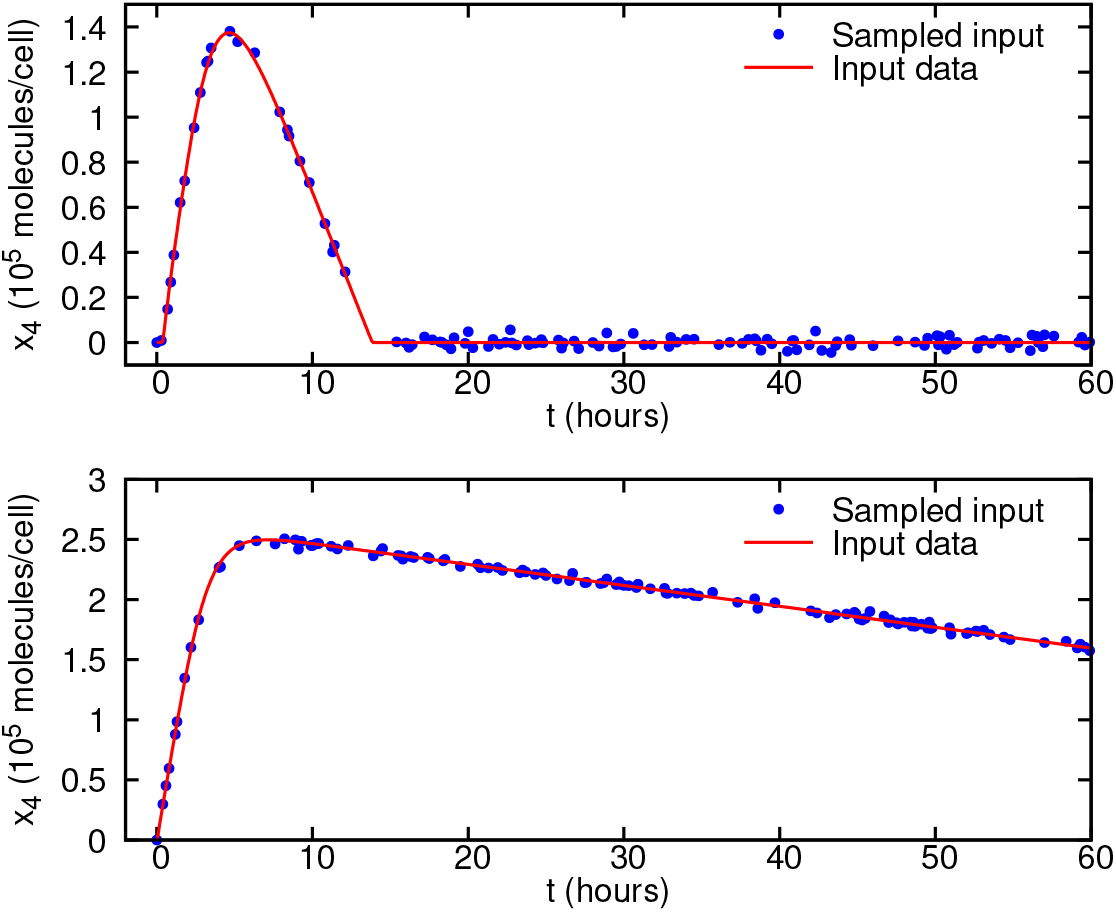
Cell apoptosis noisy observation data given to the algorithm for parameter inference. 120 measurements are corrupted by a zero-mean Gaussian noise and standard deviation of *σ* = 0.05*μ*. Data for the observable *x*_4_ only are randomly sampled during the time window of 0 – 60 hours for two scenarios: (top) cell survival with the initial condition *x*_7_(0) = 2.9 × 10^4^ (molecules/cell) and (bottom) cell death with *x*_7_(0) = 2.9 × 10^3^ (molecules/cell).

Using the systems-informed neural networks and the noisy input data, we are able to infer most of the dynamics (including *x*_3_, *x*_4_, *x*_6_, *x*_7_ and *x*_8_) of the system as shown in Figs. **S5** and **5**. These results show a good agreement between the inferred and exact dynamics of the cell survival/apoptosis models using one observable only.

**Fig. 5.**
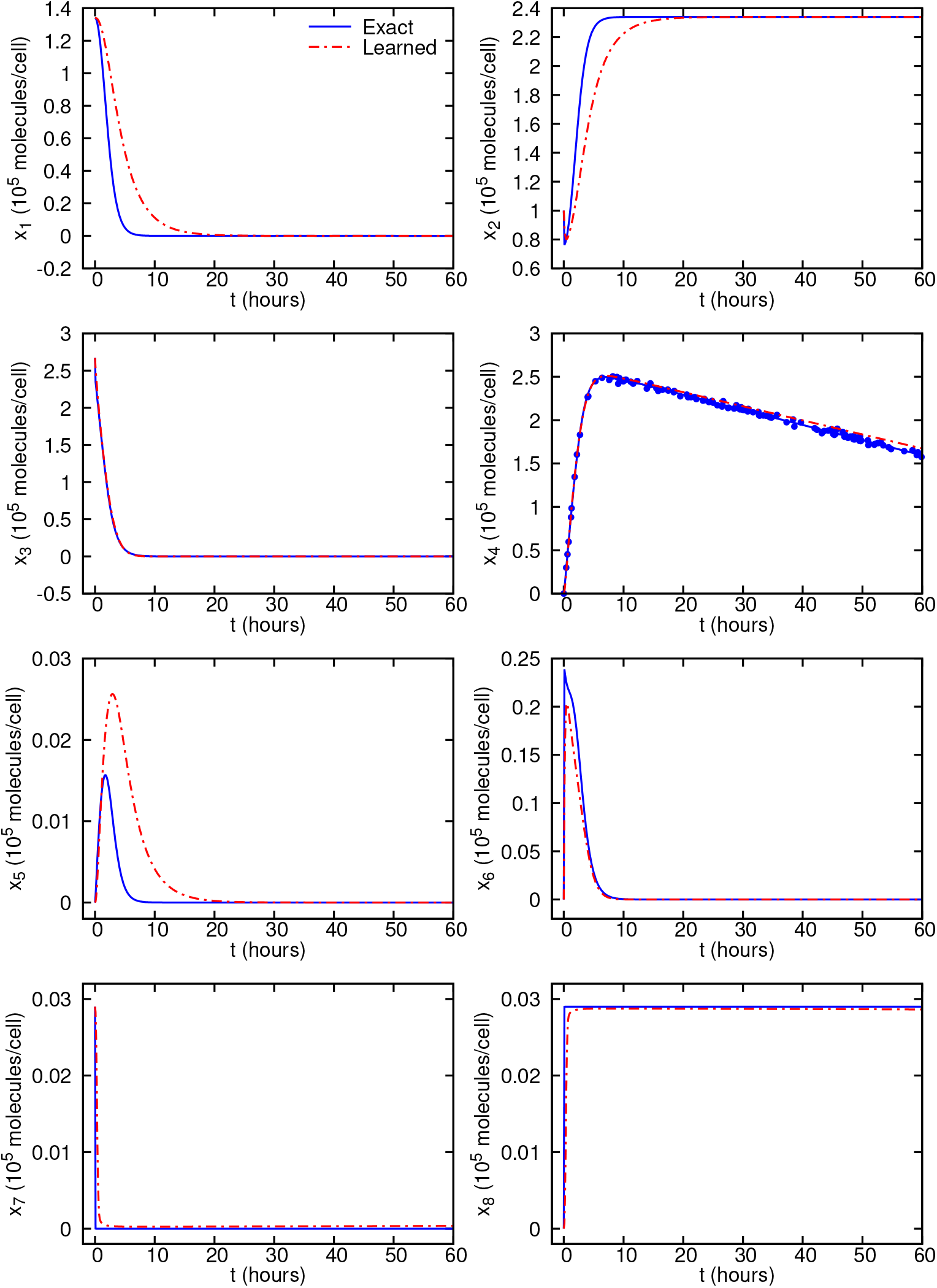
Cell apoptosis inferred dynamics from noisy observations compared with the exact solution. Predictions are performed on equally-spaced time instants in the interval of 0 – 60 hours. The scattered observations are plotted using symbols only for the observable *x*_4_. The exact data and the scattered observations are computed by solving the system of ODEs given in Eq. (S5).

We report the inferred parameters for the cell apoptosis model in Table 4, where we have used noisy observations on *x*_4_ under two scenarios of cell death and survival for comparison. The results show that four parameters (*k*_1_, *k*_*d*2_, *k*_*d*4_ and *k*_*d*6_) can be identified by our proposed method with relatively high accuracy, as indicated by the check mark (✓) in the last column of Table 4. We observe that the standard deviations for most of the parameter estimates are orders of magnitude larger than their target values, and thus the standard deviations estimated using the FIM are not informative in the practical identifiability analysis. The only informative standard deviation is for *k*_*d*6_ (indicated by the symbol †), and *k*_*d*6_ is inferred with relatively high accuracy by our method.

**Table 4.** Parameter values for cell apoptosis model and their corresponding inferred values. The standard deviations are estimated using Eq. (S3) as practical identifiability analysis using the Fisher Information Matrix. The symbols ✓, † and ✶ in the last column denote that the corresponding variable is identifiable using our proposed method, FIM standard deviation, and null-eigenvector analysis.

| Parameter | Target Value | Cell Survival |  | Cell Death |  | Identifiable |
| --- | --- | --- | --- | --- | --- | --- |
|  |  | Inferred Value | Standard Deviation | Inferred Value | Standard Deviation |  |
| $k_1$ | $2.67 \times 10^{-9}$ | $0.59 \times 10^{-9}$ | $6.2 \times 10^{-6}$ | $0.34 \times 10^{-9}$ | $1.9 \times 10^{-5}$ | $\checkmark \star$ |
| $k_{d1}$ | $1 \times 10^{-2}$ | $7.06 \times 10^{-10}$ | 35.5 | $3.37 \times 10^{-7}$ | 96.7 | |
| $k_{d2}$ | $8 \times 10^{-3}$ | $1.72 \times 10^{-3}$ | 4.9 | $2.38 \times 10^{-3}$ | 23.8 | ✓ |
| $k_3$ | $6.8 \times 10^{-8}$ | $0.15 \times 10^{-8}$ | $1.0 \times 10^{-4}$ | $0.16 \times 10^{-8}$ | $2.1 \times 10^{-4}$ | |
| $k_{d3}$ | $5 \times 10^{-2}$ | $4.19 \times 10^{-10}$ | 62.8 | $4.23 \times 10^{-10}$ | 78.0 | |
| $k_{d4}$ | $1 \times 10^{-3}$ | $0.92 \times 10^{-3}$ | 0.20 | $1.28 \times 10^{-3}$ | 1.2 | $\checkmark \star$ |
| $k_5$ | $7 \times 10^{-5}$ | $1.49 \times 10^{-7}$ | 0.019 | $7.25 \times 10^{-8}$ | 0.37 | |
| $k_{d5}$ | $1.67 \times 10^{-5}$ | $6.92 \times 10^{-11}$ | 0.034 | $8.51 \times 10^{-11}$ | 13.7 | |
| $k_{d6}$ | $1.67 \times 10^{-4}$ | $1.81 \times 10^{-4}$ | $0.35 \times 10^{-4}$ | $1.57 \times 10^{-4}$ | $1.57 \times 10^{-4}$ | $\checkmark \dagger \star$ |

To have a better picture of the practical identifiability analysis, we have plotted the correlation matrix **R** in Fig. **S6**. We observe perfect correlations |*R_ij_*| ≈ 1 between some parameters. Specifically, parameters *k*_1_ – *k*_*d*1_, and *k*_3_ – *k*_*d*3_ have correlations above 0.99 for cell survival model, which suggests that these parameters may not be identified. This is generally in agreement with the parameter inference results in Table **4** with some exceptions. Our parameter inference algorithm suggests that *k*_1_ is identifiable, whereas *k*_*d*1_ is not for the cell survival model. Thus, in order to increase the power of the practical identifiability analysis and to complement the correlation matrix, we have computed the FIM null eigenvectors and for each eigenvector we identified the most dominant coefficients, which are plotted in Fig. 6. We observe that there are six null eigenvectors associated with the zero eigenvalues of the FIM for both the cell survival and cell death models. The most dominant coefficient in each null eigenvector is associated with a parameter that can be considered as practically non-identifiable. The identifiable parameters include *k*_1_, *k*_*d*4_ and *k*_*d*6_ (indicated by the symbol ✶ in Table 4), which agree well with the results of our algorithm. On the contrary, our algorithm successfully infers one more parameter *k*_*d*2_ than the above analysis. This could be due to the fact that checking practical identifiability using the FIM can be problematic, especially for partially observed nonlinear systems [37]. We have similar results for the cell death model.

**Fig. 6.**
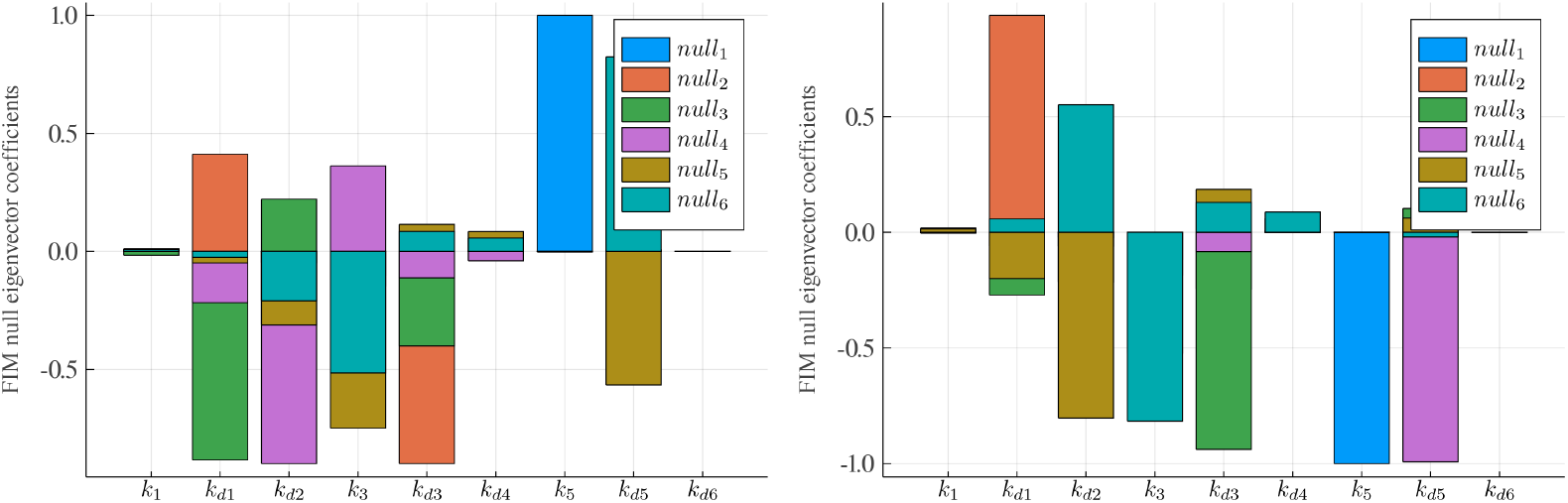
Fisher information matrix null eigenvectors of the cell apoptosis model. The most dominant component in each null eigenvector associated with a specific parameter suggests that the parameter may not be practically identifiable: (left) cell survival and (right) cell death.

### Ultradian endocrine model

The final test case for assessing the performance of the proposed algorithm is to infer parameters of the ultradian model for the glucose-insulin interaction. We use a relatively simple ultradian model [38] with 6 state variables and 30 parameters. This is a minimal model developed in a non-pathophysiologic context and represents relatively simple physiologic mechanics. In the ultradian model, the primary state variables are the glucose concentration *G*, the plasma insulin concentration *I_p_*, and the interstitial insulin concentration *I_i_*, which are appended with a three stage filter (*h*_1_, *h*_2_, *h*_3_) that reflects the response of the plasma insulin to glucose levels [38]. The resulting system of ODEs, the nominal values for the parameters of the ultradian model along with the initial conditions for the 6 state variables are given in the Supporting Information.

The nutritional driver *I_G_*(*t*) is the systematic forcing of the model that represents the external sources of glucose from nutritional intake. Although the nutritional intake (modeled by the *N* discrete nutrition events) is required to be defined and properly recorded by the patients, it is not always accurately recorded or may contain missing values. Therefore, it would be useful to employ systems-informed neural networks to not only infer the model parameters given the nutrition events, but also to assume that the intake is unknown (hidden forcing) and infer the nutritional driver in Eq. (S7)f as the same time.

#### Model parameter inference given the nutrition events

We consider an exponential decay functional form for the nutritional intake 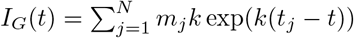, where the decay constant *k* is the only unknown parameter and three nutrition events are given by (*t_j_*, *m_j_*) = [(300, 60) (650,40) (1100, 50)] (*min, g*) pairs. The only observable is the glucose level measurements *G* shown in Fig. **7** (generated here synthetically by solving the system of ODEs), which are sampled randomly to train the neural networks for the time window of 0 – 1800 minutes. Because we only use the observation of *G* and have more than 20 parameters to infer, we limit the search range for these parameters. The range of seven parameters is adopted from [38], and the range for other parameters is set as (0.2*x*, 1.8*x*), where x is the nominal value of that parameter (Table S3).

**Fig. 7.**
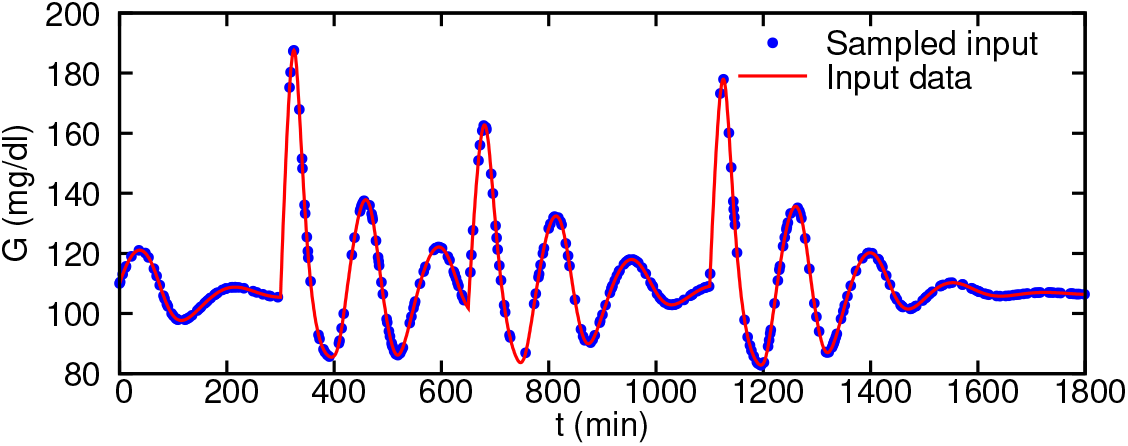
Ultradian glucose-insulin model observation data given to the algorithm for parameter inference. 360 noiseless measurements on glucose level (*G*) only are randomly sampled in the time window of 0 – 1800 minutes (~ one day).

For the first test case, we set the parameters *V_p_*, *V_i_* and *V_g_* to their nominal values and infer the rest of the parameters. The inferred values are given in Table **5** (column Test 1), where we observe good agreement between the target and inferred values. For the second test, we also infer the values of *V_p_*, *V_i_* and *V_g_* (Table 5). Although the inferred parameters are slightly worse than the Test 1, when using the inferred parameters, we are able to solve the equations for unseen time instants with high accuracy. We perform forecasting for the second test case after training the algorithm using the glucose data in the time interval of *t* = 0 – 1800 *min* and inferring the model parameters. Next, we consider that there is a nutrition event at time *t_j_* = 2000 *min* with carbohydrate intake of *m_j_* = 100 *g*. As shown in Fig. 8, we are able to forecast with high accuracy the glucose-insulin dynamics, more specifically, the glucose levels following the nutrition intake.

**Table 5.**
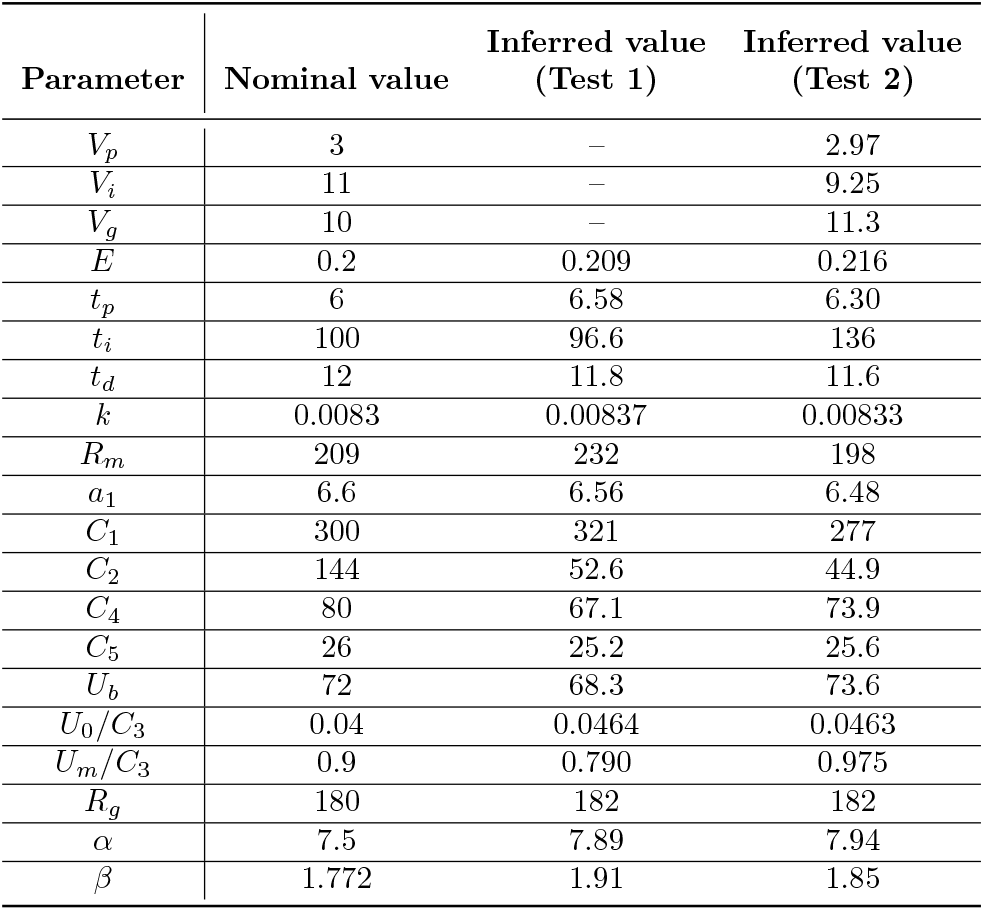
Parameter values for the ultradian glucose-insulin model and their corresponding inferred values.

| Parameter | Nominal value | Inferred value<br>(Test 1) | Inferred value<br>(Test 2) |
| --- | --- | --- | --- |
| $V_p$ | 3 | – | 2.97 |
| $V_i$ | 11 | – | 9.25 |
| $V_g$ | 10 | – | 11.3 |
| $E$ | 0.2 | 0.209 | 0.216 |
| $t_p$ | 6 | 6.58 | 6.30 |
| $t_i$ | 100 | 96.6 | 136 |
| $t_d$ | 12 | 11.8 | 11.6 |
| $k$ | 0.0083 | 0.00837 | 0.00833 |
| $R_m$ | 209 | 232 | 198 |
| $a_1$ | 6.6 | 6.56 | 6.48 |
| $C_1$ | 300 | 321 | 277 |
| $C_2$ | 144 | 52.6 | 44.9 |
| $C_4$ | 80 | 67.1 | 73.9 |
| $C_5$ | 26 | 25.2 | 25.6 |
| $U_b$ | 72 | 68.3 | 73.6 |
| $U_0/C_3$ | 0.04 | 0.0464 | 0.0463 |
| $U_m/C_3$ | 0.9 | 0.790 | 0.975 |
| $R_g$ | 180 | 182 | 182 |
| $\alpha$ | 7.5 | 7.89 | 7.94 |
| $\beta$ | 1.772 | 1.91 | 1.85 |

**Fig. 8.**
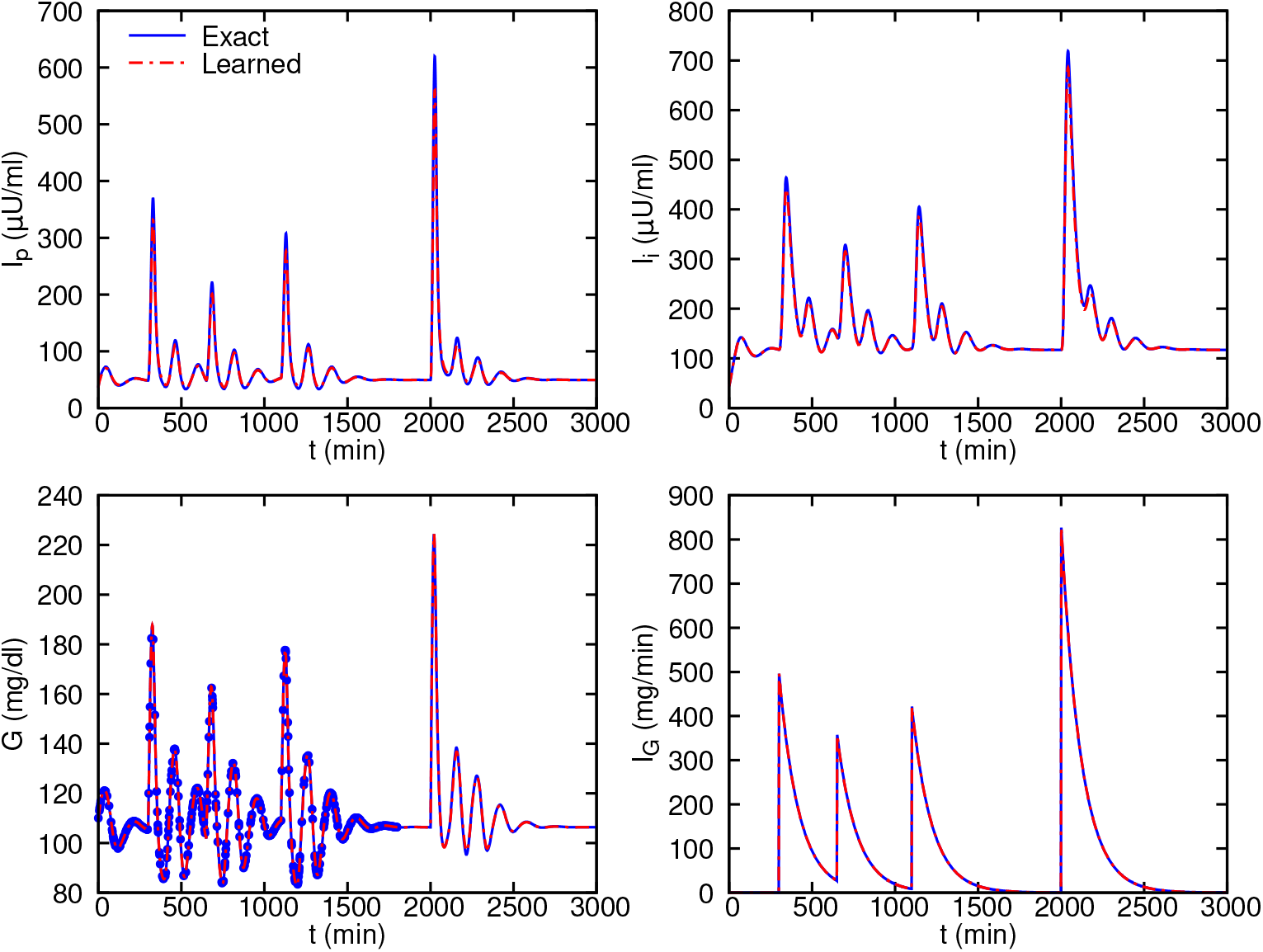
Ultradian glucose-insulin inferred dynamics and forecasting compared with the exact solution given nutrition events. 600 scattered observations of glucose level are randomly sampled from 0 – 1800 *min* and used for training. Note that the parameter *k* in the intake function *I_G_* is considered to be unknown, while the timing and carbohydrate content of each nutrition event are given. Given the inferred parameters, we can accurately forecast the glucose levels following the event at time *t* = 2000 *min*.

#### Model parameter inference with hidden nutrition events

As detailed in the following, one of the significant advantages of the systems-informed neural network is its ability to infer the *hidden* systematic forcing in the model. For example, in the glucose-insulin model, the nutritional driver *I_G_* is the forcing that we aim to infer as well. Here, we use the glucose measurements to train the model for the time interval *t* = 0 – 1800 *min* shown in Fig. 7, while we assume that the time instants and quantities of three nutritional events are additionally unknown.

We found that it is difficult to infer all the parameters as well as the the timing and carbohydrate content of each nutrition event. However, given *V_p_*, *V_i_*, *V_g_* and the timing of each nutrition event, the algorithm is capable of inferring the other model parameters as well as the carbohydrate content (Table **S4** column Test 1). Having the nutrition events as well as all other unknown parameters estimated, we are able to forecast the glucose levels for *t* = 1800 – 3000 *min* assuming there has been a nutritional intake of (*t_j_*, *m_j_*) = (2000,100). The predictions for the glucose *G* and the nutritional driver *I_G_* are shown in Fig. S7, which show excellent agreement in the forecasting of glucose levels. For the second test, we also infer the values of *V_p_*, *V_i_* and *V_g_*, and the result is slightly worse (Table **S4** column Test 2 and Fig. S8).

If both the timing and carbohydrate content of each nutrition event are unknown, the algorithm is also capable to infer them by assuming that certain model parameters are known. We found that the selection of the known parameters is important. As shown in Table S4, we consider different combinations of parameters to be known in Test 3 and Test 4; Test 3 leads to good prediction accuracy (Fig. S9) while Test 4 does not.

## Discussion

We presented a new and simple to implement “systems-biology-informed” deep learning algorithm that can reliably and accurately infer the *hidden* dynamics described by a mathematical model in the form of a system of ODEs. The system of ODEs is encoded into a plain “uninformed” deep neural networks and is enforced through minimizing the loss function that includes the residuals of the ODEs. Enforcing the equations in the loss function adds additional constraints in the learning process, which leads to several advantages of the proposed algorithm: first, we are able to infer the unknown parameters of the system of ODEs once the neural network is trained; second, we can use a minimalistic amount of data on a few observables to infer the dynamics and the unknown parameters; third, the enforcement of the equations adds a regularization effect that makes the algorithm robust to noise (we have not used any other regularization technique); and lastly, the measurements can be scattered, noisy and just a few.

The problem of structural and practical non-identifiability (such as the one encountered in the cell apoptosis model) is a long-standing problem in the field of systems identification, and has been under extensive research, e.g., [39]. Structural non-identifiabilities originate from incomplete observation of the internal model states. Since a structural non-identifiability is independent of the accuracy of available experimental data, it cannot be resolved by a refinement of existing measurements, and one way to resolve this issue is through increasing the number of observed species. Our focus was mostly on practical identifiability, which can guide us to redesign the experiment, modify the model or collect additional measurements. In this study, we used FIM and local sensitivity analysis for the identifiability analysis, but we note that FIM has many limitations and can be problematic, especially for partially observed nonlinear systems [37], and hence other advanced alternatives [15, 40] should be used in future works. However, our goal in this work was not to do systematic identifiability analysis, but rather to use identifiability analysis to explain some of our findings.

## Conclusion

We have used three benchmark problems to assess the performance of the algorithm including a highly nonlinear glycolysis model, a non-identifiable cell apoptosis model, and an ultradian glucose-insulin model for glucose forecasting based on the nutritional intake. Given the system of ODEs and initial conditions of the state variables, the algorithm is capable of accurately inferring the whole dynamics with one or two observables, where the unknown parameters are also inferred during the training process. An important and very useful outcome of the algorithm is its ability to infer the systematic forcing or driver in the model such as the nutritional intake in the glucose-insulin model. In this work, we considered the synthetic data of three small problems to test the performance and limitation of the proposed method. We will apply our method to larger problems and real data (e.g., the dataset introduced in [41]) in future work.

## Supporting information

**S1 Text. Supporting information for “Systems biology informed deep learning for inferring parameters and hidden dynamics”.**

## Acknowledgments

This work was supported by the National Institutes of Health U01HL142518. Computations were supported by the National Science Foundation XSEDE resources award No. TG-DMS140007.

