## Supplementary Information for "Systems biology informed deep learning for inferring parameters and hidden dynamics"

### 1 Fisher information matrix

The Fisher information matrix (FIM) is constructed by the following equation and by estimating the local sensitivities of the system of ODEs given in Eq. (1) with respect to the parameters

$$\mathbf{F} = \sum_{t_n} \mathbf{S}^T \cdot \mathbf{C}^{-1} \cdot \mathbf{S}, \tag{S1}$$

where $\mathbf{C}$ is the covariance matrix of the measurements error and $\mathbf{S} = \partial \mathbf{x} / \partial \mathbf{p}$ is the sensitivity matrix (partial derivatives of the state variables with respect to the parameters). Because the state variables are time-dependent, the sensitivities are also time-dependent. Thus, a set of differential equations in the following form has to be solved for $\mathbf{S}_n$ at each time instant $t_n$:

$$\dot{\mathbf{S}} = \frac{\partial \boldsymbol{f}}{\partial \mathbf{x}} \cdot \mathbf{S} + \frac{\partial \boldsymbol{f}}{\partial \mathbf{p}}, \tag{S2}$$

where $\mathbf{J} \equiv \partial \boldsymbol{f} / \partial \mathbf{x}$ is the Jacobian of the system. FIM is then calculated according to Eq. (S1) by summing up all values over the time span. Note that FIM is the inverse of the parameter estimation error covariance matrix of the best linear unbiased estimator. The standard deviations of the parameter estimates are therefore the squared roots of the diagonal elements of $\mathbf{F}^{-1}$. They are, however, only lower bounds for the standard deviations as the system is nonlinear in the parameters [1]. Thus, we can use the standard deviations to estimate 95% confidence intervals of the parameter estimates using the following equations:

$$\begin{aligned} p_i - 2\sigma_i \leq p_i^* \leq p_i + 2\sigma_i, \\ \sqrt{F_{ii}^{-1}} \leq \sigma_i, \end{aligned} \tag{S3}$$

where $p_i^*$ is the "true" value of the parameter $p_i$.

It is also possible to check the practical identifiability using FIM-based criteria, where a (nearly) singular FIM indicates non-identifiable parameters due to non-informative experiments (assuming the model is structurally identifiable). The high correlations among parameters may lead to a singular FIM. Thus, we investigate this using two approaches: we construct the correlation matrix $R_{ij} = F_{ij}^{-1} / F_{ii}^{-1}$ and then search for correlations of $|R_{ij}| \approx 1.0$ between the parameters; and we compute the eigenvetors of FIM associated with the zero eigenvalues (i.e., *null* eigenvectors) to look for non-identifiable parameters. Specifically, a null eigenvector having a dominant component associated with a single parameter indicates that changes in this parameter do not affect the state variables. The local sensitivity analysis is generally not able to detect structurally non-identifiable parameters. Therefore, our analysis based on FIM is primarily focused on practical identifiability.

### 2 Yeast glycolysis model

The equations for the yeast glycolysis model are given in Eq. (S4). To generate synthetic data, the system of ODEs is solved using the solver `odeint` of `SciPy` library from time $t = 0$ to $t = 10\ min$ with the following initial conditions: $\mathbf{S}(0) = [0.501\ 1.955\ 0.198\ 0.148\ 0.161\ 0.161\ 0.064]$ (mM).

The model consists of ODEs for the concentrations of seven biochemical species:

$$\frac{dS_1}{dt} = J_0 - \frac{k_1 S_1 S_6}{1+(S_6/K_1)^q}, \tag{S4a}$$

$$\frac{dS_2}{dt} = 2\frac{k_1 S_1 S_6}{1+(S_6/K_1)^q} - k_2 S_2 (N - S_5) - k_6 S_2 S_5, \tag{S4b}$$

$$\frac{dS_3}{dt} = k_2 S_2 (N - S_5) - k_3 S_3 (A - S_6), \tag{S4c}$$

$$\frac{dS_4}{dt} = k_3 S_3 (A - S_6) - k_4 S_4 S_5 - \kappa (S_4 - S_7), \tag{S4d}$$

$$\frac{dS_5}{dt} = k_2 S_2 (N - S_5) - k_4 S_4 S_5 - k_6 S_2 S_5, \tag{S4e}$$

$$\frac{dS_6}{dt} = -2\frac{k_1 S_1 S_6}{1+(S_6/K_1)^q} + 2k_3 S_3 (A - S_6) - k_5 S_6, \tag{S4f}$$

$$\frac{dS_7}{dt} = \psi\kappa (S_4 - S_7) - k S_7, \tag{S4g}$$

where the parameters for the model are taken from [2] and listed in Table **S1**.

### 3 Cell apoptosis model

A system of ODEs describing the activation of $x_3$ by $x_1$ is constructed using mass-action kinetics and the known topology of the pathways and is given by Eq. (S5). The parameters of the system are given in Table **S2**. The system of ODEs is solved using `odeint` from time $t = 0$ to $t = 60$ *hours* with the initial conditions given as $\mathbf{x}(0) = [1.34 \times 10^5\ 1.0 \times 10^5\ 2.67 \times 10^5\ 0.0\ 0.0\ 0.0\ x_7^0\ 0.0]$ $(molecules/cell)$, where $x_7^0 = 2.9 \times 10^3$ leads to cell death and $x_7^0 = 2.9 \times 10^4$ leads to cell survival. It should be mentioned that a non-dimensional form of the Eq. (S5) is encoded into the neural networks loss function, where the network is trained with non-dimensional observation on $x_4$. To non-dimensionalize the data, a timescale of $t_{scale} = 3600\ s$ and concentration scale of $c_{scale} = 1.0 \times 10^5\ molecules/cell$ are used.

The equations defining the cell apoptosis model include eight species and are as follows:

$$\frac{dx_1}{dt} = -k_1x_4x_1 + k_{d1}x_5, \quad \text{(S5a)}$$

$$\frac{dx_2}{dt} = k_{d2}x_5 - k_3x_2x_3 + k_{d3}x_6 + k_{d4}x_6, \quad \text{(S5b)}$$

$$\frac{dx_3}{dt} = -k_3x_2x_3 + k_{d3}x_6, \quad \text{(S5c)}$$

$$\frac{dx_4}{dt} = k_{d4}x_6 - k_1x_4x_1 + k_{d1}x_5 - k_5x_7x_4 + k_{d5}x_8 + k_{d2}x_5, \quad \text{(S5d)}$$

$$\frac{dx_5}{dt} = -k_{d2}x_5 + k_1x_4x_1 - k_{d1}x_5, \quad \text{(S5e)}$$

$$\frac{dx_6}{dt} = -k_{d4}x_6 + k_3x_2x_3 - k_{d3}x_6, \quad \text{(S5f)}$$

$$\frac{dx_7}{dt} = -k_5x_7x_4 + k_{d5}x_8 + k_{d6}x_8, \quad \text{(S5g)}$$

$$\frac{dx_8}{dt} = k_5x_7x_4 - k_{d5}x_8 - k_{d6}x_8, \quad \text{(S5h)}$$

where the values of the rate constants for the model are taken from [3] and listed in Table **S2**.

### 4 Ultradian endocrine model

The system of equations of the glucose-insulin interaction model is given by Eqs. (S6) and (S7) for which the major parameters include: (i) $E$, a rate constant for exchange of insulin between the plasma and remote compartments; (ii) $I_G$, the exogenous (externally driven) glucose delivery rate; (iii) $t_p$, the time constant for plasma insulin degradation; (iv) $t_i$, the time constant for the remote insulin degradation; (v) $t_d$, the delay time between plasma insulin and glucose production; (vi) $V_p$, the volume of insulin distribution in the plasma; (vii) $V_i$, the volume of the remote insulin compartment; (viii) $V_g$, the volume of the glucose space [4, 5]. Nominal values of the model parameters are given in Table **S3**. Further, in Eq. (S7), $f_1(G)$ represents the rate of insulin production; $f_2(G)$ represents insulin-independent glucose utilization; $f_3(I_i)$ represents insulin-dependent glucose utilization; $f_4(h_3)$ represents delayed insulin-dependent glucose utilization.

$$\frac{dI_p}{dt} = f_1(G) - E\left(\frac{I_p}{V_p} - \frac{I_i}{V_i}\right) - \frac{I_p}{t_p}, \tag{S6a}$$

$$\frac{dI_i}{dt} = E\left(\frac{I_p}{V_p} - \frac{I_i}{V_i}\right) - \frac{I_i}{t_i}, \tag{S6b}$$

$$\frac{dG}{dt} = f_4(h_3) + I_G(t) - f_2(G) - f_3(I_i)G, \tag{S6c}$$

$$\frac{dh_1}{dt} = \frac{1}{t_d}(I_p - h_1), \tag{S6d}$$

$$\frac{dh_2}{dt} = \frac{1}{t_d}(h_1 - h_2), \tag{S6e}$$

$$\frac{dh_3}{dt} = \frac{1}{t_d}(h_2 - h_3), \tag{S6f}$$

where $f_1 - f_4$ and the nutritional driver of the model $I_G(t)$ are given by

$$f_1(G) = \frac{R_m}{1 + \exp(\frac{-G}{V_g c_1} + a_1)}, \tag{S7a}$$

$$f_2(G) = U_b\left(1 - \exp(\frac{-G}{C_2 V_g})\right), \tag{S7b}$$

$$f_3(I_i) = \frac{1}{C_3 V_g}\left(U_0 + \frac{U_m}{1 + (\kappa I_i)^{-\beta}}\right), \tag{S7c}$$

$$f_4(h_3) = \frac{R_g}{1 + \exp(\alpha(\frac{h_3}{C_5 V_p} - 1))}, \tag{S7d}$$

$$\kappa = \frac{1}{C_4}\left(\frac{1}{V_i} + \frac{1}{E t_i}\right), \tag{S7e}$$

$$I_G(t) = \sum_{j=1}^{N} m_j k \exp(k(t_j - t)), \tag{S7f}$$

where $I_G(t)$ is defined over $N$ discrete nutrition events [6] with $k$ as the decay constant and event $j$ occurs at time $t_j$ with carbohydrate quantity $m_j$.

The nutritional driver of the model is the intake $I_G(t)$ defined over $N$ discrete nutritional events given in Eq. (S7)f. To generate the synthetic data for training, the system of ODEs is solved using `odeint` from time $t = 0$ to $t = 1800$ $min$ with the initial conditions given as $\mathbf{x}(0) = [12.0\ (\mu U/ml)\ 4.0\ (\mu U/ml)\ 110.0\ (mg/dl)\ 0.0\ 0.0\ 0.0]$ and three nutrition events given by $(t_j, m_j) = [(300, 60)\ (650, 40)\ (1100, 50)]\ (min, g)$ pairs.

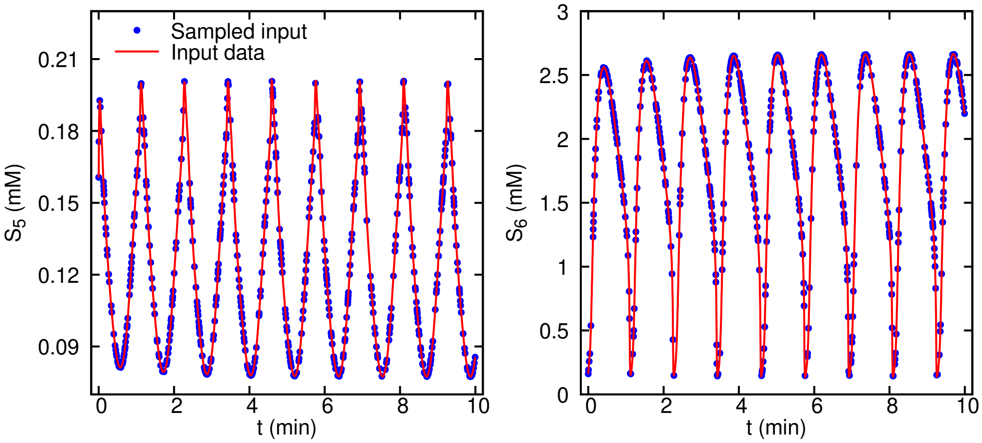


**Fig. S1. Glycolysis oscillator noiseless observation data given to the algorithm for parameter inference.** 500 noiseless measurements of two observables $S_5$ and $S_6$ are randomly sampled in the time window of $0 - 10$ minutes.

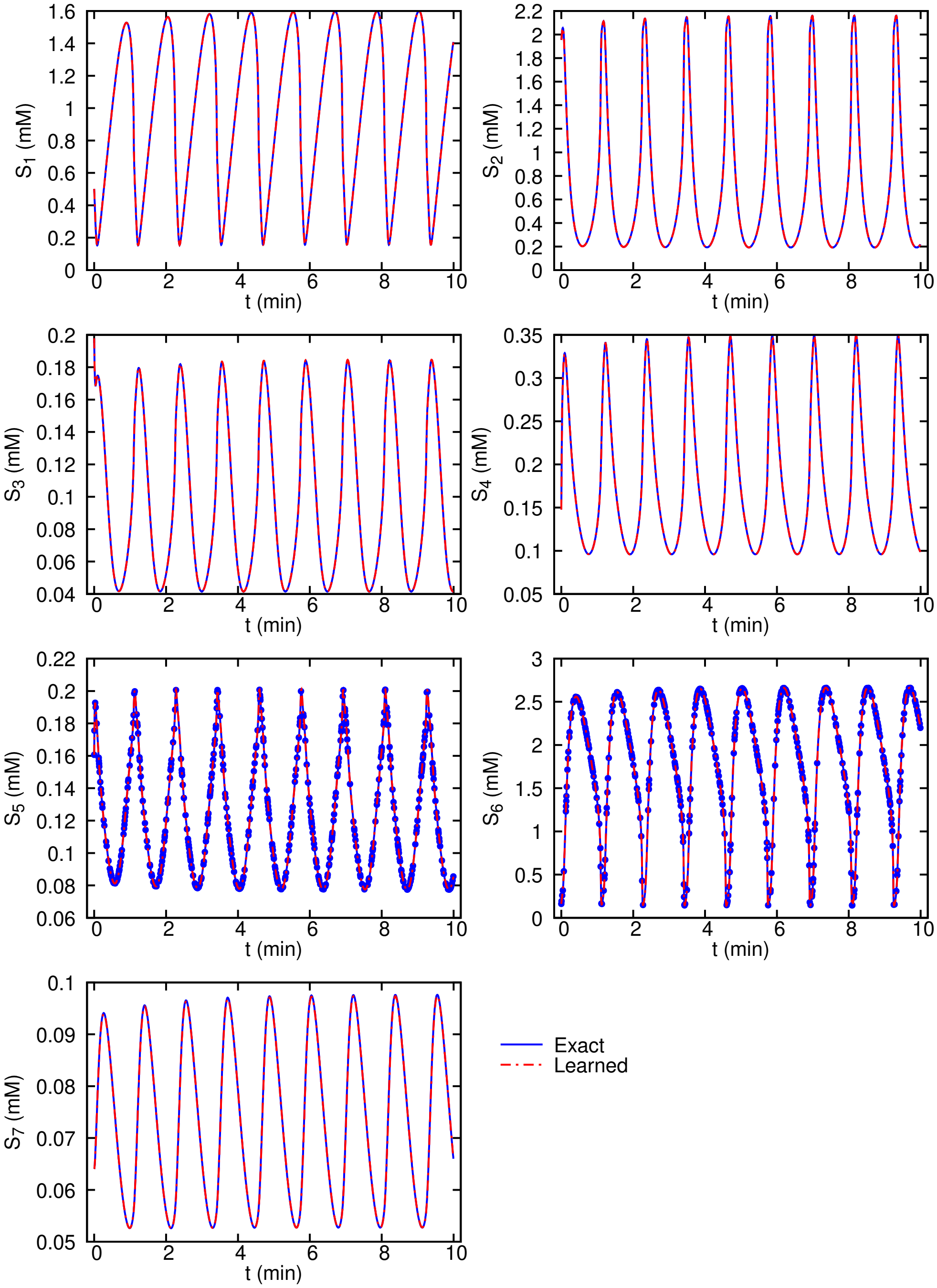


**Fig. S2. Glycolysis oscillator inferred dynamics compared with the exact solution.** Predictions are performed on equally-spaced time instants in the interval of $0-10$ minutes. The scattered observations are plotted using symbols for the two observables $S_5$ and $S_6$. The exact data and the scattered observations are computed by solving the system of ODEs given in Eq. (S4).

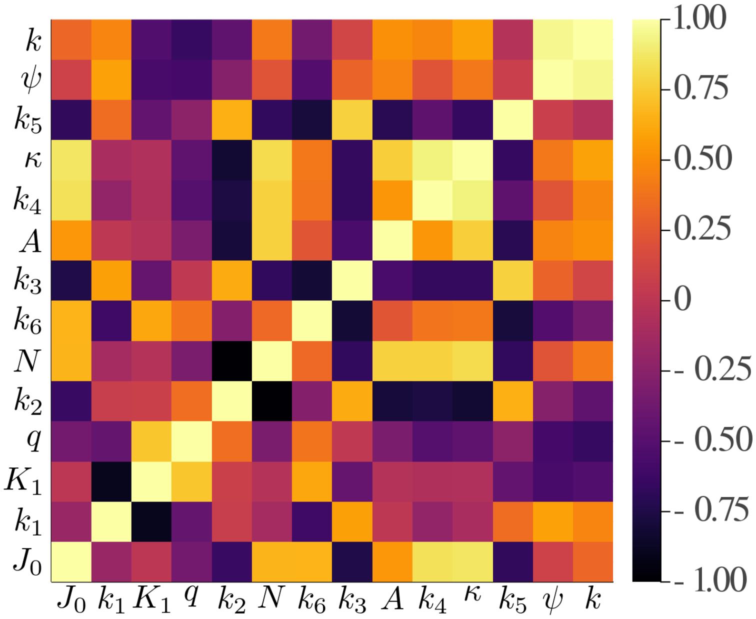


**Fig. S3. Correlation matrix for the parameters of glycolysis model.** The correlation matrix is computed using the local sensitivity analysis and the FIM for the parameters involved in the glycolysis oscillator model assuming 10% noise in the observation data. We observe no perfect correlations suggesting that FIM is not singular and the parameters in the glycolysis model are practically identifiable.

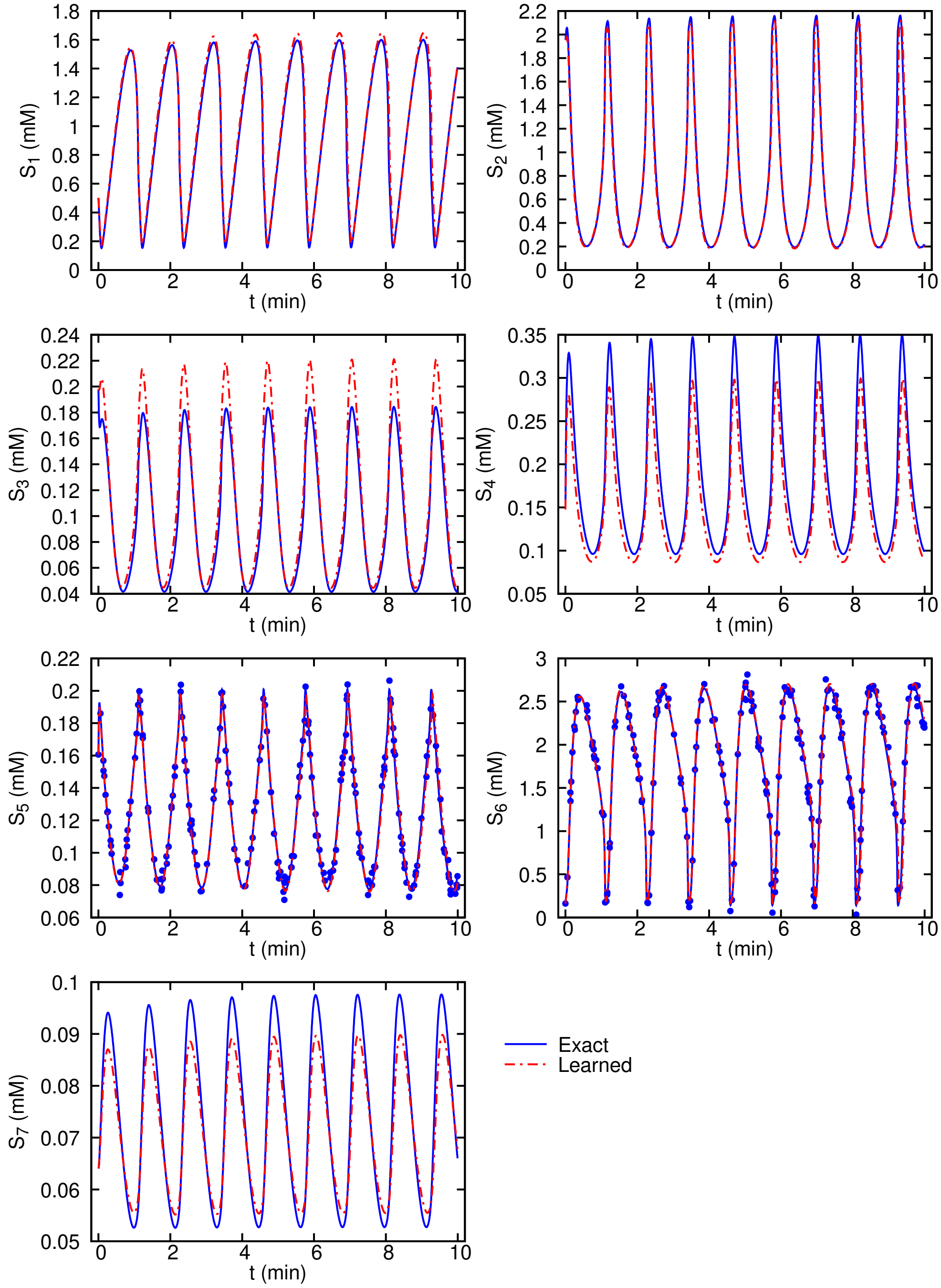


**Fig. S4. Glycolysis oscillator inferred dynamics from noisy measurements compared with the exact solution.** 200 scattered observations are plotted using symbols for the two observables $S_5$ and $S_6$.

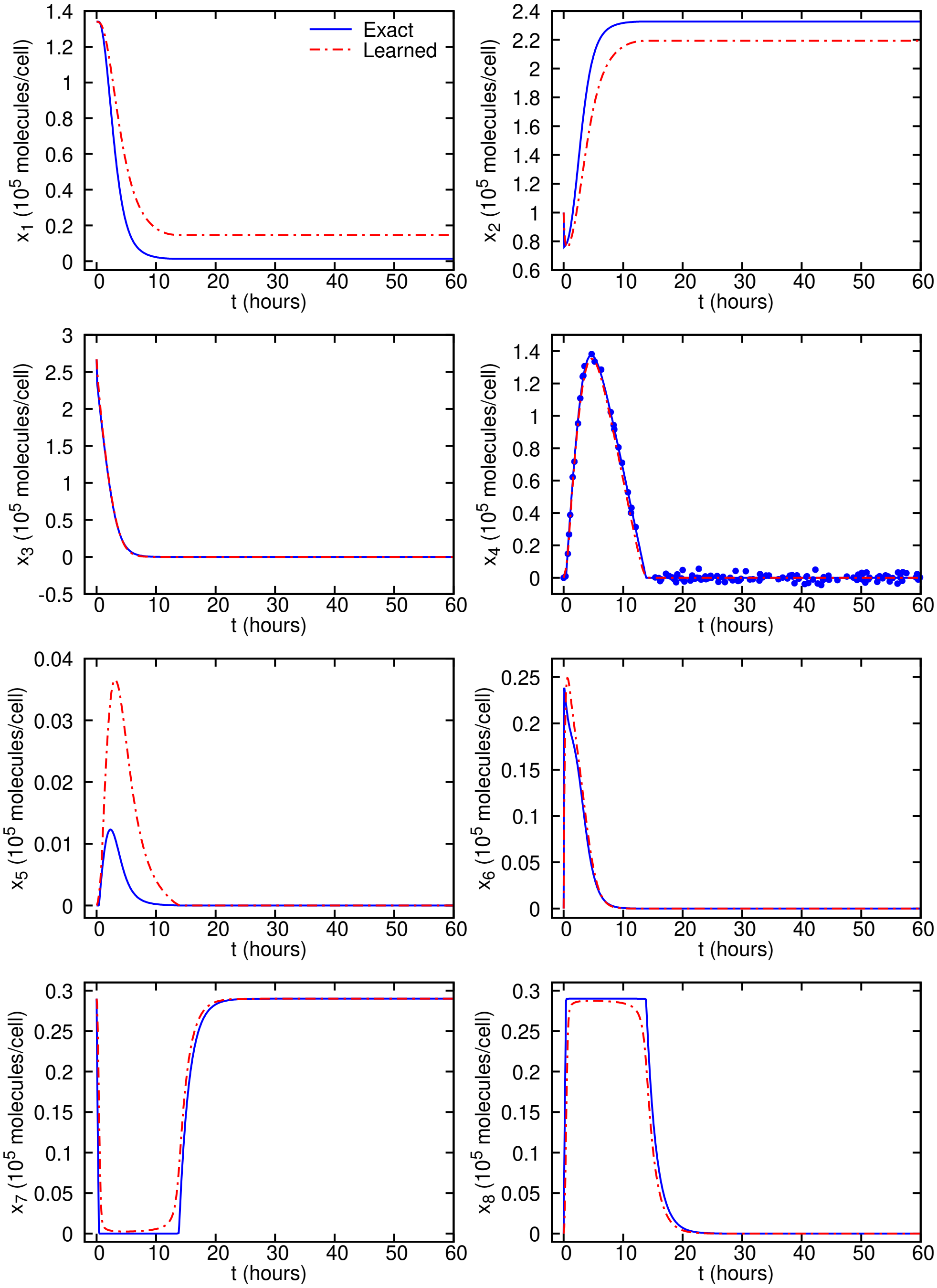


**Fig. S5. Cell survival inferred dynamics from noisy observations compared with the exact solution.** Predictions are performed on equally-spaced time instants in the interval of $0 - 60$ hours. The scattered observations are plotted using symbols only for the observable $x_4$. The exact data and the scattered observations are computed by solving the system of ODEs given in Eq. (S5).

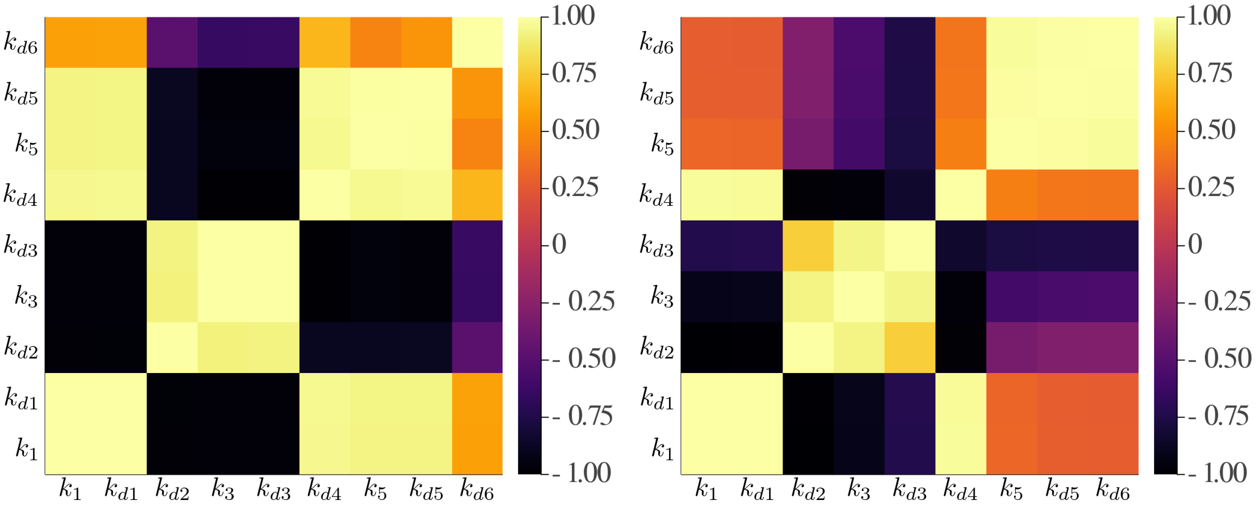


**Fig. S6. Correlation matrix for the parameters of the cell apoptosis model.** The correlation matrix is computed using FIM for the practical identifiablity analysis of parameters involved in the cell apoptosis model assuming 5% noise in the observation data for two scenarios: (left) cell survival and (right) cell death. We observe perfect correlations of $\approx 1.0$ between the parameters suggesting that the FIM is singular and some parameters in the cell apoptosis model are practically non-identifiable.

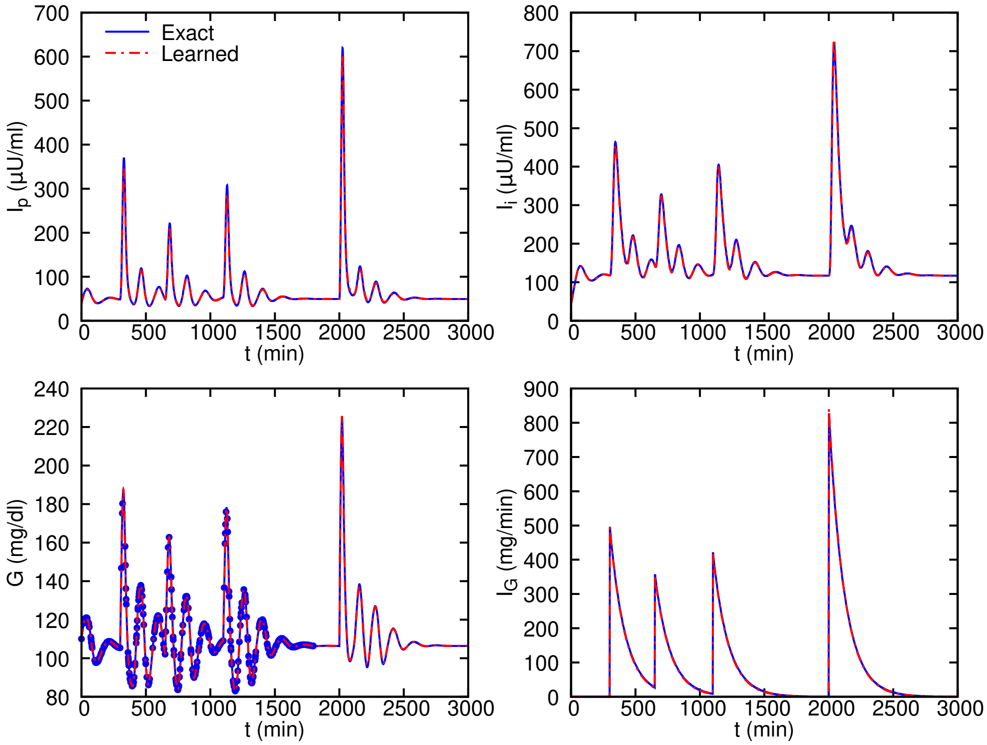


**Fig. S7. Ultradian glucose-insulin inferred dynamics with hidden nutritional driver (Test 1 in Table S4).** Scattered observations of glucose level are randomly sampled from $0 - 1800$ $min$ and used for training. The parameter $k$ in the intake function $I_G$ as well as carbohydrate content $(m_j)$ of each nutrition event are treated as unknown, while $V_p, V_i, V_g$ and the timing $(t_j)$ of each nutrition event are given.

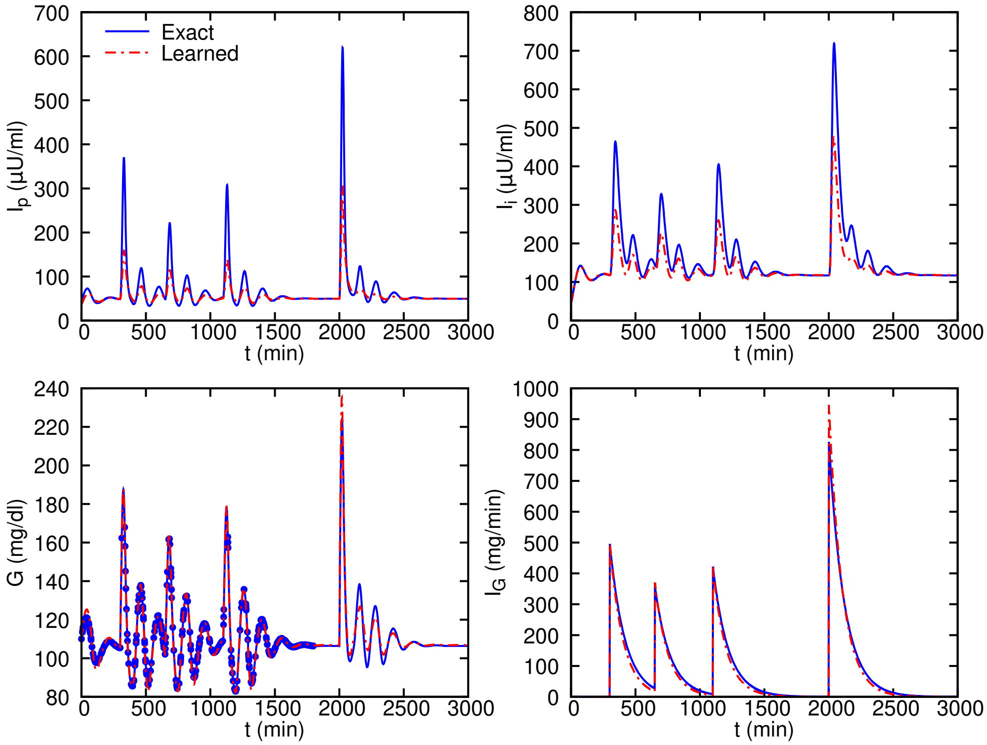


**Fig. S8. Ultradian glucose-insulin inferred dynamics with hidden nutritional driver (Test 2 in Table S4).** Scattered observations of glucose level are randomly sampled from $0 - 1800$ $min$ and used for training. The parameter $k$ in the intake function $I_G$ as well as carbohydrate content $(m_j)$ of each nutrition event are treated as unknown, while the timing $(t_j)$ of each nutrition event are given.

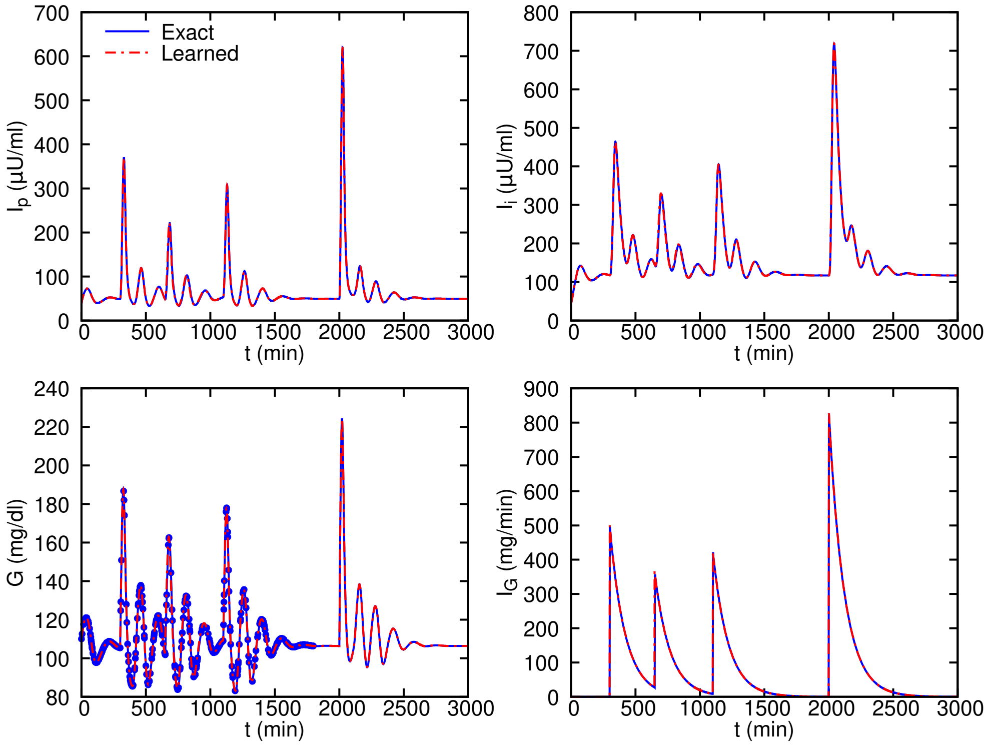


**Fig. S9. Ultradian glucose-insulin inferred dynamics with hidden nutritional driver (Test 3 in Table S4).** Scattered observations of glucose level are randomly sampled from $0 - 1800$ $min$ and used for training. The timing $(t_j)$ and carbohydrate content $(m_j)$ of each nutrition event are treated as unknown, while the parameter $k$ in the intake function $I_G$ is given.

**Table S1. Full list of parameters for glycolytic oscillator model [2].**

| Parameter | Nominal value | Unit |
|---|---|---|
| $J_0$ | 2.5 | $mM\ min^{-1}$ |
| $k_1$ | 100 | $mM^{-1}\ min^{-1}$ |
| $k_2$ | 6 | $mM^{-1}\ min^{-1}$ |
| $k_3$ | 16 | $mM^{-1}\ min^{-1}$ |
| $k_4$ | 100 | $mM^{-1}\ min^{-1}$ |
| $k_5$ | 1.28 | $min^{-1}$ |
| $k_6$ | 12 | $mM^{-1}\ min^{-1}$ |
| $k$ | 1.8 | $min^{-1}$ |
| $\kappa$ | 13 | $min^{-1}$ |
| $q$ | 4 | |
| $K_1$ | 0.52 | $mM$ |
| $\psi$ | 0.1 | |
| $N$ | 1 | $mM$ |
| $A$ | 4 | $mM$ |

**Table S2. Full list of parameters for cell apoptosis model [3].**

| Parameter | Nominal value | Unit |
|---|---|---|
| $k_1$ | $2.67 \times 10^{-9}$ | cell $\cdot$ ($s$ $\cdot$ molecules)$^{-1}$ |
| $k_{d1}$ | $1 \times 10^{-2}$ | $s^{-1}$ |
| $k_{d2}$ | $8 \times 10^{-3}$ | $s^{-1}$ |
| $k_3$ | $6.8 \times 10^{-8}$ | cell $\cdot$ ($s$ $\cdot$ molecules)$^{-1}$ |
| $k_{d3}$ | $5 \times 10^{-2}$ | $s^{-1}$ |
| $k_{d4}$ | $1 \times 10^{-3}$ | $s^{-1}$ |
| $k_5$ | $7 \times 10^{-5}$ | cell $\cdot$ ($s$ $\cdot$ molecules)$^{-1}$ |
| $k_{d5}$ | $1.67 \times 10^{-5}$ | $s^{-1}$ |
| $k_{d6}$ | $1.67 \times 10^{-4}$ | $s^{-1}$ |

**Table S3. Full list of parameters for the ultradian glucose-insulin model [5].** The search range for the first 7 parameters is adopted from [4], and the range for the other parameters is $(0.2x, 1.8x)$, where $x$ is the nominal value of that parameter.

| Parameter | Nominal value | Unit | Search range |
|---|---|---|---|
| $V_p$ | 3 | $lit$ | (2, 4) |
| $V_i$ | 11 | $lit$ | (7, 15) |
| $V_g$ | 10 | $lit$ | (7, 13) |
| $E$ | 0.2 | $lit\ min^{-1}$ | (0.1, 0.3) |
| $t_p$ | 6 | $min$ | (4, 8) |
| $t_i$ | 100 | $min$ | (60, 140) |
| $t_d$ | 12 | $min$ | (25/3, 50/3) |
| $k$ | 0.0083 | $min^{-1}$ | |
| $R_m$ | 209 | $mU\ min^{-1}$ | |
| $a_1$ | 6.6 | | |
| $C_1$ | 300 | $mg\ lit^{-1}$ | |
| $C_2$ | 144 | $mg\ lit^{-1}$ | |
| $C_3$ | 100 | $mg\ lit^{-1}$ | |
| $C_4$ | 80 | $mU\ lit^{-1}$ | |
| $C_5$ | 26 | $mU\ lit^{-1}$ | |
| $U_b$ | 72 | $mg\ min^{-1}$ | |
| $U_0$ | 4 | $mg\ min^{-1}$ | |
| $U_m$ | 90 | $mg\ min^{-1}$ | |
| $R_g$ | 180 | $mg\ min^{-1}$ | |
| $\alpha$ | 7.5 | | |
| $\beta$ | 1.772 | | |

**Table S4. Parameter values for the ultradian glucose-insulin model with hidden nutritional driver and their corresponding inferred values.**

| Parameter | Nominal value | Inferred value (Test 1) | Inferred value (Test 2) | Inferred value (Test 3) | Inferred value (Test 4) |
|---|---|---|---|---|---|
| $t_j$ | 300, 650, 1100 | – | – | 299.8, 647.5, 1100 | 294, 640, 1093 |
| $m_j$ | 60, 40, 50 | 58.8, 39.3, 49.0 | 52.3, 37.3, 43.8 | 60.1, 40.8, 50.1 | 33.4, 21.9, 21.1 |
| $V_p$ | 3 | – | 2.52 | – | – |
| $V_i$ | 11 | – | 7.01 | – | – |
| $V_g$ | 10 | – | 12.3 | – | – |
| $E$ | 0.2 | 0.219 | 0.300 | – | – |
| $t_p$ | 6 | 6.74 | 8.00 | 6.00 | – |
| $t_i$ | 100 | 91.8 | 140 | 99.8 | – |
| $t_d$ | 12 | 11.7 | 10.7 | 12.0 | – |
| $k$ | 0.0083 | 0.00849 | 0.00955 | – | – |
| $R_m$ | 209 | 206 | 376 | – | 41.8 |
| $a_1$ | 6.6 | 6.54 | 6.02 | – | 3.22 |
| $C_1$ | 300 | 312 | 423 | – | 540 |
| $C_2$ | 144 | 51.4 | 42.9 | 74.2 | 259 |
| $C_4$ | 80 | 73.5 | 45.4 | 78.7 | 42.2 |
| $C_5$ | 26 | 25.5 | 24.9 | 26.0 | 15.8 |
| $U_b$ | 72 | 72.8 | 93.5 | 71.3 | 14.4 |
| $U_0/C_3$ | 0.04 | 0.0394 | 0.0288 | 0.0399 | 0.008 |
| $U_m/C_3$ | 0.9 | 0.837 | 0.823 | 0.886 | 1.62 |
| $R_g$ | 180 | 180 | 189 | 179 | 324 |
| $\alpha$ | 7.5 | 7.78 | 13.5 | 7.51 | 13.5 |
| $\beta$ | 1.772 | 1.82 | 2.86 | 1.78 | 3.19 |
